# A genetic network coordinated by *TCP16* and *LHY* integrates regulation of the vegetative-reproductive phase transition in *Arabidopsis thaliana*

**DOI:** 10.64898/2026.05.26.727858

**Authors:** Pouya Motienoparvar, Ali Ebrahimi, Kaveh Kavousi, Mokhtar Jalali Javaran, Charles Spillane, Peter C. McKeown

## Abstract

The transition to flowering in *Arabidopsis thaliana* is a complex process governed by many biological and environmental stimuli. Although many of the genes which regulate this process have been identified over the past 30 years, it remains unclear how these networks are integrated. In this study, we used the transcriptional responses of Col-0, L*er*-1, and three mutant lines, to build a genome wide regulatory network of *Arabidopsis thaliana* during the flowering transition. The expression profiles of 22,810 genes across five genotypes were collected from the GEO database Series GSE57 from which we assigned flowering-time genes to different interacting modules by an adapted form of Hierarchical Complete Linkage Clustering (HCLC) after reconstruction of regulatory networks according to the Position Weight Matrix (PWM)-based method. Within these modules, we identified 77 ‘core’ genes and 31 controller or ‘driver’ genes. We identify two genes, *LHY* and, less expectedly, the transcription factor *TCP16*, to be topographically positioned at the regulatory hubs a nine-gene transcriptional control unit, implying they have the capacity to integrate information from across the flowering time pathways which interpret different environmental or endogenous cues during the vegetative-reproductive transition. Interrogating their behaviour across transcriptional datasets, we show that both *LHY* and *TCP16* show transcriptional oscillations during the flowering transition, with a wavelength that varies depending on environmental conditions. We suggest that the transcriptional responses of *LHY* and *TCP16* allow them to regulate the flow of information through the genetic networks which integrates different floral transition cues, and that genetic modelling approaches can provide new insights into the regulation of well-studied biological processes such as the flowering transition.

**Author summary:** How plants decide when to flower is a critical stage for completing their life cycles. It is also of key agricultural importance, as crops need to flower at the right time of year to allow efficient pollination and harvesting. Many genes are known to affect flowering time control in plants. Here, we use computational approaches to estimate how different genes interact in flowering time control in Arabidopsis, a small plant in the mustard family which is widely used for molecular studies. We use large-scale studies of how gene expression changes in different plant lines which have disrupted or adjusted flowering time to group the many genes involved in flowering into different interacting pathway, which we visualise as sets of coloured ‘nodes’ controlling one another in a network. We show that two genes may have new rols in integrating information from different pathways, and discuss how their behaviour might help them to function as intregrators of biological information – including the daily oscaillations in their expression.

## Introduction

One of the most important phenological characteristics during the domestication and breeding of plants during the history of agriculture is the timing of flower emergence. The transition to flowering can be understood as an example of chronobiology, in which the plant’s capacity to measure time through circadian rhythms, intrinsic mechanisms and environmental cues is translating into growth and development outcomes. By better manipulating pathways underlying plant chronobiology, breeders may be enabled to optimize yield, improve responses to environmental stresses (such as by drought avoidance), deploy speed breeding, and ultimately, create more resilient and productive plant varieties [1,2]. Humans have optimized the flowering time to be able to grow plants in the desired environments [3]. In recent decades, many studies have been conducted to identify genes involved in controlling the timing of the vegetative to reproductive phase transition, but more precise management of flowering time remains challenging. In fact, studies have shown that environmental factors as diverse as day length, light intensity, drought, temperature and various soil characteristics can combine with plant genotype to orchestrate the phenology of flowering [1, 4–7].

The vegetative to reproductive phase transition in the model eudicot *Arabidopsis thaliana* (L.) Heynh (Brassicaceae), hereafter Arabidopsis, is regulated by a complex network of genes and signaling pathways that respond to both endogenous and environmental cues [8–10]. Arabidopsis thaliana is a facultative long-day (LD) plant, meaning it exhibits accelerated expression of the FT (*FLOWERING LOCUS T*) gene under LD conditions. LD-dependent flowering is associated with the activation of the photoperiodic pathway through the transcriptional regulator CO (CONSTANS), which belongs to the B-box (BBX) Zn-finger family. The transition requires the activation of a group of genes called floral meristem identity (FMI) genes which promote the formation of floral meristems. The FMI genes are regulated by many upstream regulators, including the photoperiod pathway, the vernalization pathway, and the autonomous pathway. Together these regulate the expression of a core group of genes, *FLOWERING LOCUS C* (*FLC*), *FLOWERING LOCUS T* (*FT*) and *SUPPRESSOR OF OVER EXPRESSION OF CONSTANS* (*SOC1*), which collectively control the meristem transition itself (Figure 1; reviewed [11]). The photoperiod pathway is activated by changes in day length and involves the regulation of *CONSTANS*, which promotes the expression of the FMI genes [12–15]. The vernalization pathway is activated by exposure to cold temperatures and involves the regulation of the *FLOWERING LOCUS C* (*FLC*) gene, which represses the expression of the *FMI* genes [16]. *FLC* functions as a central inhibitor of flowering and is the primary determinant of the winter cold requirement for flowering [17–18]. In winter annuals, *FLC* is activated by the FRIGIDA (FRI) complex (FRI, FLC EXPRESSOR (FLX), FLX-LIKE 4 (FLX4)), with FLX and FLX4 also playing essential roles in establishing basal *FLC* expression in summer annuals [19–22].

**Figure 1.**
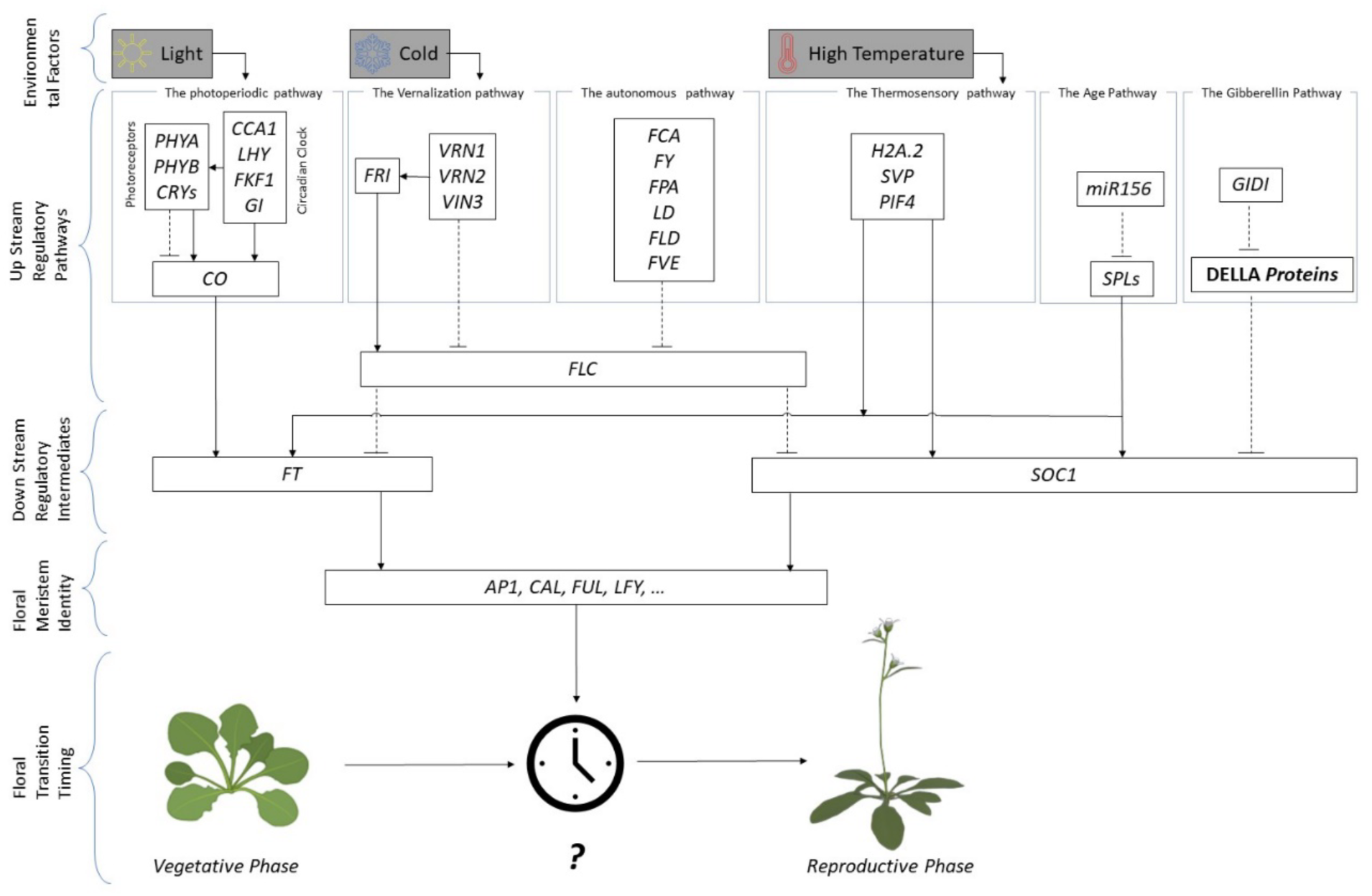
Simplified genetic pathways regulating flowering in *Arabidopsis thaliana*. Environmental and endogenous inputs are shown at the top, leading through regulatory pathways to the floral meristem identity (FMI) genes leading to control of floral transition timing; six pathways and a subset off their component genes are enclosed in the dashed boxes; lines with arrows indicate activation, and broken lines with blunt ends indicate repression. How these signals are integrated is indicated by the interrogation mark at center bottom. Modified from Banta and Heyland (2011).

The autonomous pathway is instead independent of environmental cues and involves the regulation of genes such as *FLOWERING LOCUS D* (*FLD*) and *FLOWERING LOCUS K* (*FLK*), which promote FMI gene expression [23–24]. The transition is also affected by many other genes and signaling pathways, including the gibberellin, cytokinin pathway, and miRNA pathways [25–26].

Roles have also been reported for mechanisms as divergent as DNA methylation [27], histone modification enzymes [28], proline synthesis [29] and the nonsense-mediated mRNA decay (NMD) surveillance system [30]. These pathways interact to form a complex regulatory network that ultimately controls the timing and progression of the transition.

While many of the biochemical and genetic interactions between genes (and gene products) involved in the flowering transition have been determined, these approaches do not necessarily capture how information flows through, and is integrated by, biological systems. In contrast, genetic network modelling allows entire the responses of entire systems to be determined, allowing better integration of experimental results and more accurate predictions of phenotypic responses [31–33]. A genetic network can be considered a form of data processing network [34–36] and the modelling approach taken to develop them depends on the aspects of the regulatory network prioritized and the level of detail desired. For instance, Boolean networks provide a coarse-grained view that is valuable for understanding qualitative dynamics of gene expression changes [37–38] but cannot capture continuous changes or stochastic fluctuations in gene expression. Continuous dynamical models, such as differential equations, offer a more detailed view over time and can be used for quantitative analysis of regulatory interactions [39]. Stochastic models incorporate the inherent randomness and noise of biological systems and are useful for understanding the variability shown by the flowering transition. Bayesian networks supply a probabilistic way of inferring regulatory relationships from experimental data [40], whereas information theory-based models can quantify the information flow within a network. These models can be combined to provide complementary information [41]. Graphical Gaussian models can also reveal the conditional dependences between genes, also helping to identify the key regulators [42].

A full model of flowering control will require accurate maps of the interactomes between transcription factors and their targets, and robust genome-wide transcriptomes derived under different environmental conditions to allow their dynamics to be reliably tested. Models of the genetic regulatory networks which control Arabidopsis flowering have been developed using some of these approaches, albeit without integrating statistical tests of robustness. For example, Wang and colleagues (2014) created a dynamic model of flowering time regulation in Arabidopsis considering light as an environmental factor using the S-system, the Michaelis-Menten model and the Mass-action model as their main approaches [43]. However, the existence of multiple genetic parameters, the inability to test the robustness of the resulting network statistically, and the use of only environmental parameters mean the resulting model is highly simplified. Similarly, Leal Valentim and colleagues used expression courses of eight genes known *a priori* to be involved in flowering time (*SHORT VEGETATIVE PHASE (SVP)*, *FLOWERING LOCUS C (FLC)*, *AGAMOUS-LIKE 24 (AGL24)*, *SUPPRESSOR OF OVEREXPRESSION OF CONSTANS 1 (SOC1)*, *APETALA1 (AP1)*, *FLOWERING LOCUS T (FT)*, *LEAFY (LFY)* and *FD*.to determine a dynamic network of key regulators, identifying unexpected levels of control between them [8].

In the present study, we extend these approaches by using an information theory-based modeling approach informed by whole-genome transcriptomes to identify the smallest possible numbers of integrator genes whose expression determines the length of the plant’s vegetative period. We identify the interactions between genes which respond to the flowering transition, and the regulatory modules to which they belong, centred upon a data processing unit which canalizes and integrates the data flow from other units and which we term the *[vegetative-reproductive] transition control unit*. Finally, we examine the behavior of key genes within this unit across the transition from vegetative to reproductive stage to suggest new insights into how the flowering transition is controlled.

## Results

### Combined impact of Genotype, temperature, and light Intensity on Flowering Time

In this study, we investigated the flowering time of six *Arabidopsis thaliana* genotypes: Col-0, Ler, *lfy-12* (in the Col-0 background), *co-2* (in the Ler background), and *ft-2* (also in the Ler background), under varying environmental conditions. Plants were grown under short-day conditions (9 hours light/15 hours dark) for 15 days and then transferred to long-day conditions (16 hours light/8 hours dark). The experiment included two light intensities (140 µmol m⁻² s⁻¹ and 80 µmol m⁻² s⁻¹) and three temperatures (15 °C, 23 °C, and 28 °C). For the wild type Col-0, the effect of temperature under control light intensity (100%) was monitored every day and phenological events were recorded (Table 1).

**Table 1.**
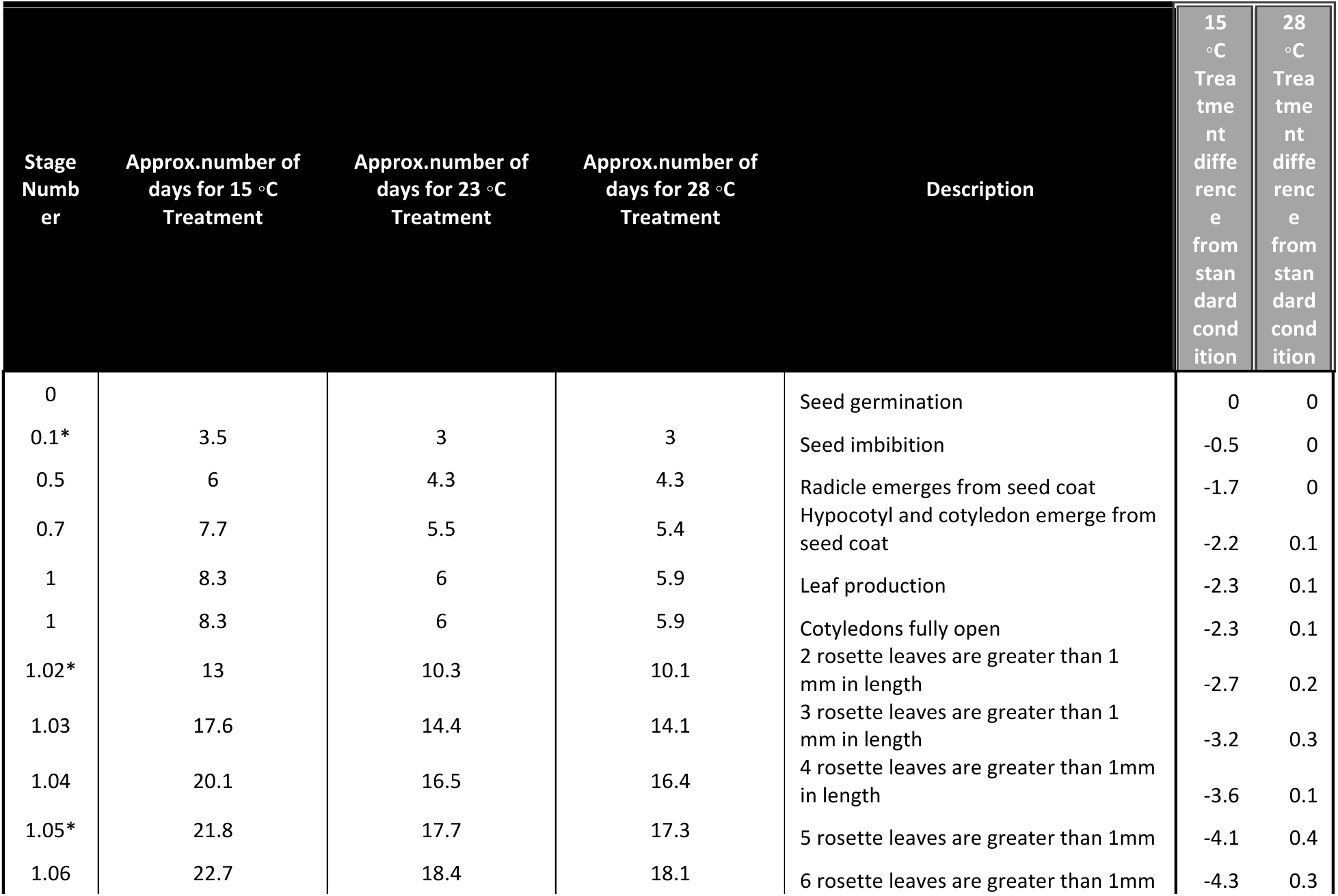
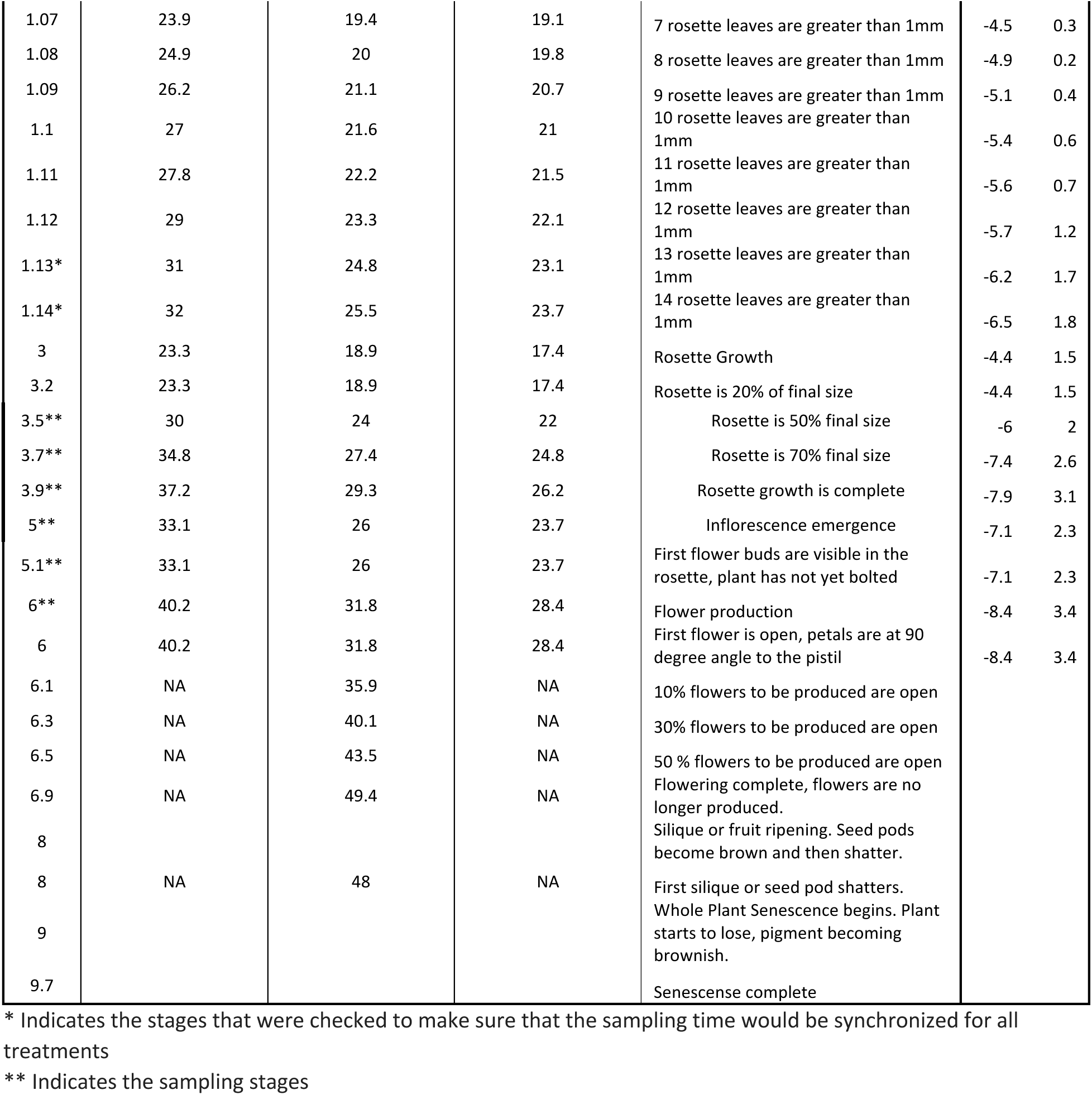
Growth stages of *Arabidopsis thaliana* Col-0 genotype in three temperature conditions including 15, 23 and 28 °C. Number of days that took to see the mentioned biological stage is listed, as are the differences from standard conditions (final two columns).

The flowering time for the Col-0 genotype varied significantly with temperature and light intensity, as expected. At 15 °C, flowering was delayed, with Col-0 plants flowering between 40 and 45 days post-germination under the higher light intensity (140 µmol m⁻² s⁻¹) and 45 to 50 days under the lower light intensity (80 µmol m⁻² s⁻¹). The highest temperature (28 °C) resulted in the earliest flowering, with plants flowering between 28 and 33 days under high light intensity and 33 to 38 days under low light intensity. Similar trends were observed for the Ler genotype, with earlier flowering at higher temperatures and light intensities (Data S3). However, Ler plants consistently flowered earlier than Col-0 under all conditions, reflecting known genotype differences [44].

The *lfy-12* mutant, which disrupts the *LEAFY* gene, exhibited a significant delay in flowering across all environmental conditions. In most cases, flowering was delayed indefinitely, and some plants failed to flower entirely, regardless of temperature or light intensity. This result is consistent with previous studies that have shown that LEAFY is critical for floral meristem identity and transition from vegetative to reproductive growth [45]. The severe delay in flowering in the *lfy-12* mutant underlines the crucial role of LEAFY in promoting flowering under long-day conditions. Similarly, the *co-2* mutant, which affects the *CONSTANS* gene, showed delayed flowering compared to wild-type Ler under all tested conditions. At 15 °C and 80 µmol m⁻² s⁻¹, *co-2* plants flowered between 50 and 55 days, which was the most delayed flowering observed for this mutant. In contrast, at 28 °C and 140 µmol m⁻² s⁻¹, flowering occurred between 35 and 40 days. These findings also align with the established role of CONSTANS in promoting flowering under long-day conditions, where loss-of-function mutations like *co-2* result in significant flowering delays [46]. The *ft-2* mutant, affecting the *FLOWERING LOCUS T* (*FT*) gene, exhibited the most severe delays in flowering, with many plants failing to flower across all conditions. When flowering did occur, it was highly delayed, taking over 50 days post-germination, as expected for this key integrator of various flowering pathways [47–48]. These results emphasize the necessity of FT for timely flowering, even under favorable long-day conditions and confirm the expected phenotypes of the germplasm used for subsequent development of the co-expression networks.

### Weighted Gene Co-expression analysis of the Arabidopsis flowering control network

To determine the network dynamics of the genes which control the flowering transition in Arabidopsis (Figure 1) and identify the modules and submodules which integrate different forms of information (as shown schematically in Figure 2), we compiled a time-course of genome-wide transcriptional responses of five genotypes to being transitioned from short-day to long-day conditions, across two replicates. The samples used were two wild-type accessions, Col-0 and L*er*-1, and the well-characterized flowering time mutants *lfy-12* (in the Col-0 background) and *co-2* and *ft-2* (both in the L*er*-1 background). Normalized expression data was determined for each of the five genotypes (Figure 3) and the relationships between genes and genotypes characterized as heatmaps (Figure 3A). Clusters representing the two genetic backgrounds, Col-0 and L*er*-1, could both be easily distinguished by visual examination (Figure 3B). Notably, the experimental replicates appear to clearly cluster together, suggesting that the design of the analysis was robust.

**Figure 2.**
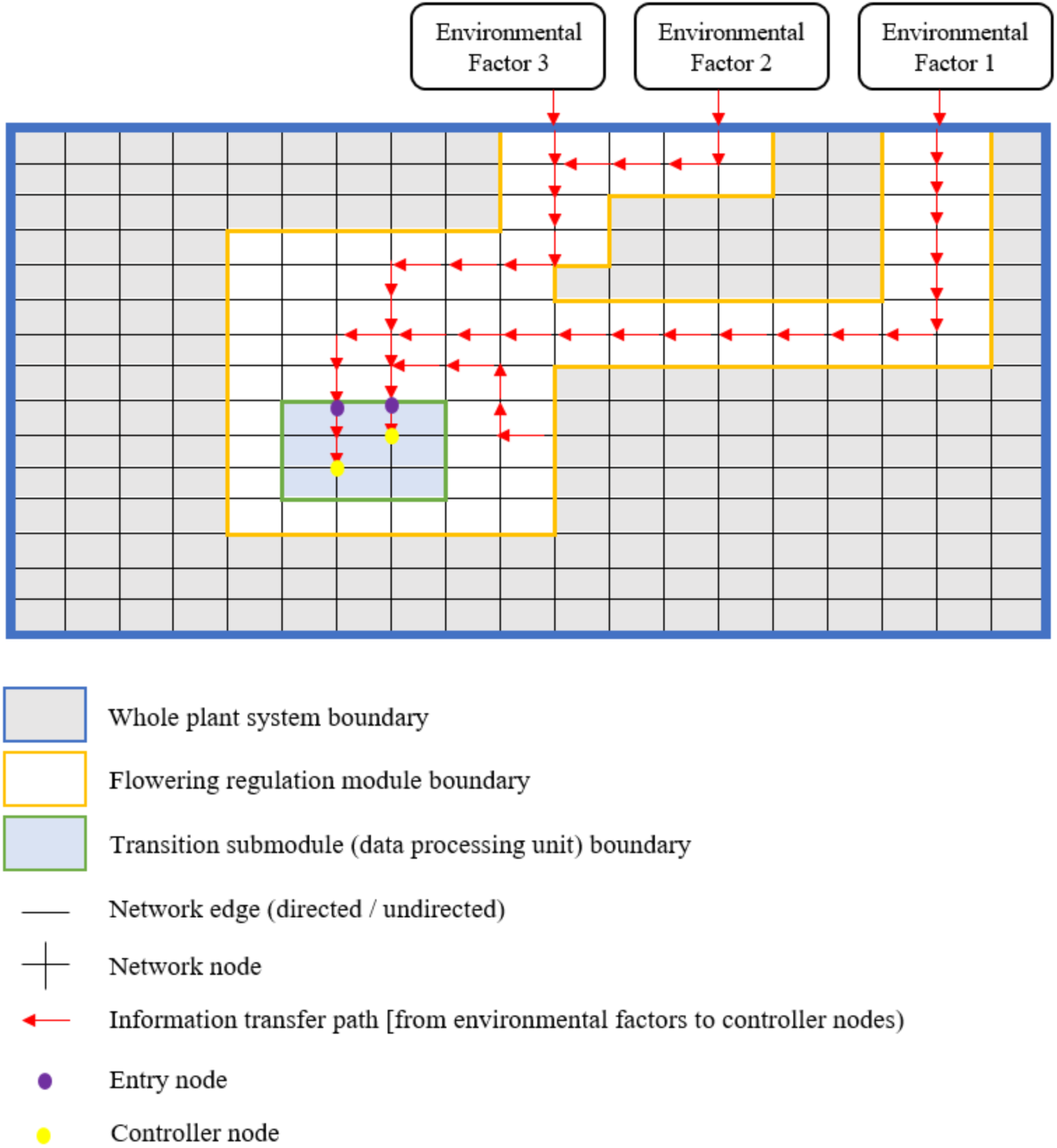
Schematic representation of a systemic approach to understanding the flow of information in the control of flowering time. It is assumed that environmental factors and genetic components of Arabidopsis form a complex system containing dedicated subsystems (modules) which control of specific biological processes (e.g. flowering). Inside these subsystems are smaller networks that regulate the features related to the flowering process, e.g. the timing of transition to reproductive phase (as shown in Figure 1). Although many nodes, edges and paths can enter or leave a node, not all necessarily carry information that has significant phenological effects; the information path can eventually lead to the controlling nodes, whose quality of expression has a great impact on the control of flowering time. Input nodes are the nodes through which information enters the [transition] network and reaches the controller nodes.

**Figure 3.**
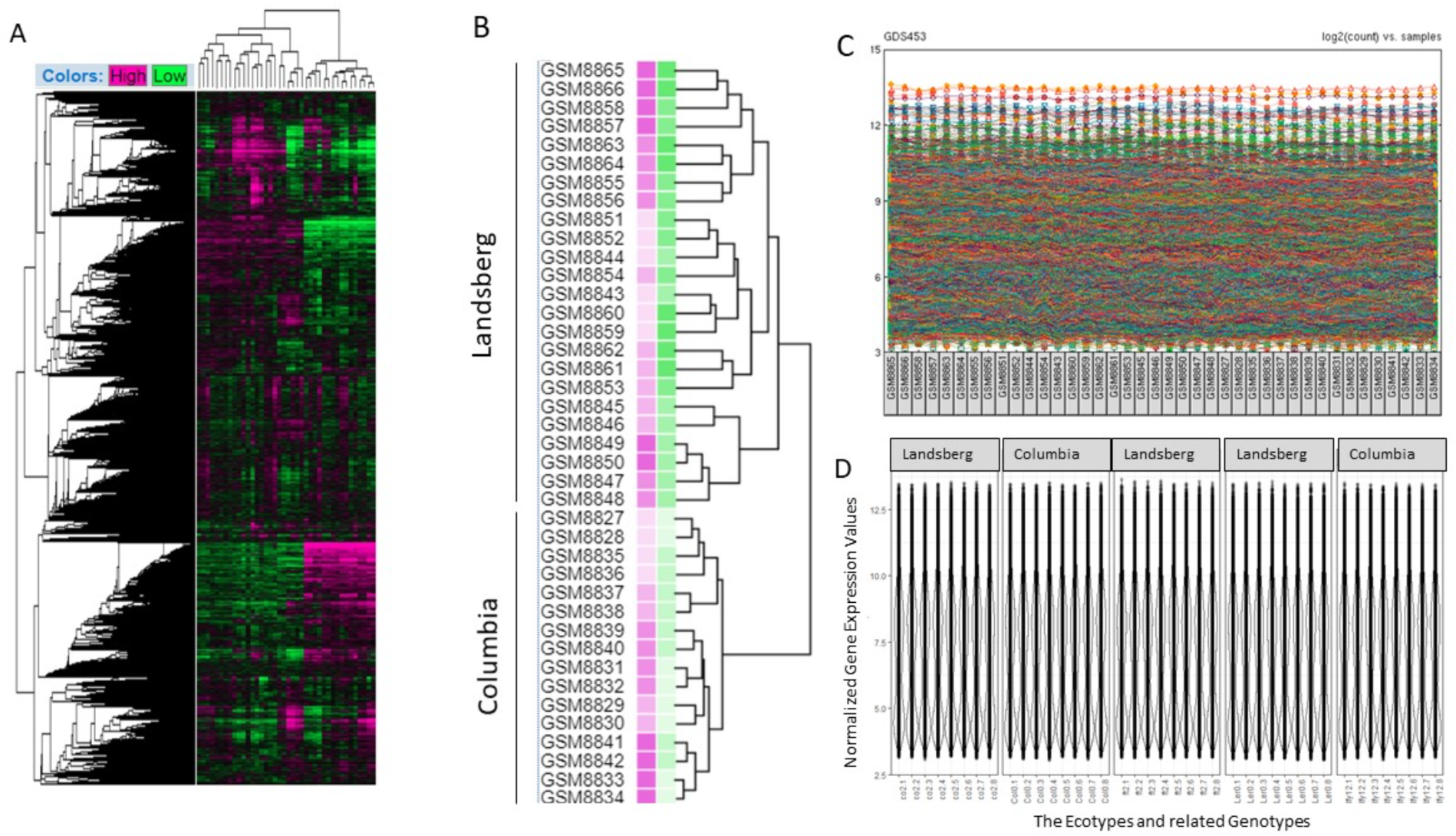
An overview of normalized expression data. (A), heatmap of the relationship between genes (left) and genotypes (top) illustrating gene expression levels in each genotype. (B) disaggregation of gene expression in the two wild-type genotypes, Columbia and Landsberg; individual samples are easily discernible as is the grouping of replicates. (C, D), assessment of the standardization of gene expression by sample (C) and by genotype (D).

This analysis was used to explore the subnetwork structures governing the flowering process. Adopting a systems approach, we displayed the connections between genes in the form of nodes and edges belonging to an information system or network, including all the physiological changes that must be coordinated during the transition to flowering process. Initially, data integrity was upheld through stringent assessments of normality, and the correlation landscape between genes and genotypes was scrutinized. To confirm that our method correctly resolved identified flowering time genes as such, we selected 46 genes that had been previously determined to affect the flowering process (Data S1) and analyzed their distribution in the identified modules. Pearson correlation was used to capture linear associations between expression profiles, aligning with the assumption of a linear relationship inherent in the method. Subsequently, we expanded the module by incorporating genes exhibiting high Pearson correlation coefficients (90% of the highest correlations) with the initial seed genes. We then applied Weighted Gene Co-expression Network Analysis (WGCNA) to characterize the co-expression networks governing flowering and validated the reproducibility of the experimental replicates using a Welch Two Sample t-test which gave p=0.7387 (>0.05), allowing us to retain the null hypothesis, i.e. that the means of Replicate 1 and Replicate 2 are statistically equal, as suggested by the clustering.

To ensure the appropriateness of the soft thresholding power (β), we leveraged the scale-free topology model fit, selecting the power of 5 through an examination of mean connectivity (Figure 4). Subsequently, the modules which comprise this network were identified, allowing us to categorize all 22,810 genes whose expression levels had been interrogated across the time-course in the five genotypes into 32 distinct modules. These were distinguished by arbitrary attribution of a color (Figure 5). The largest module, *turquoise*, encompassed a remarkable 2,978 genes, approximately 13% of the total set. At the other end of the spectrum, the smallest module, *paleturquoise*, comprised a modest 64 genes, or 0.28% of the total gene pool (Figure 5A, C).

**Figure 4.**
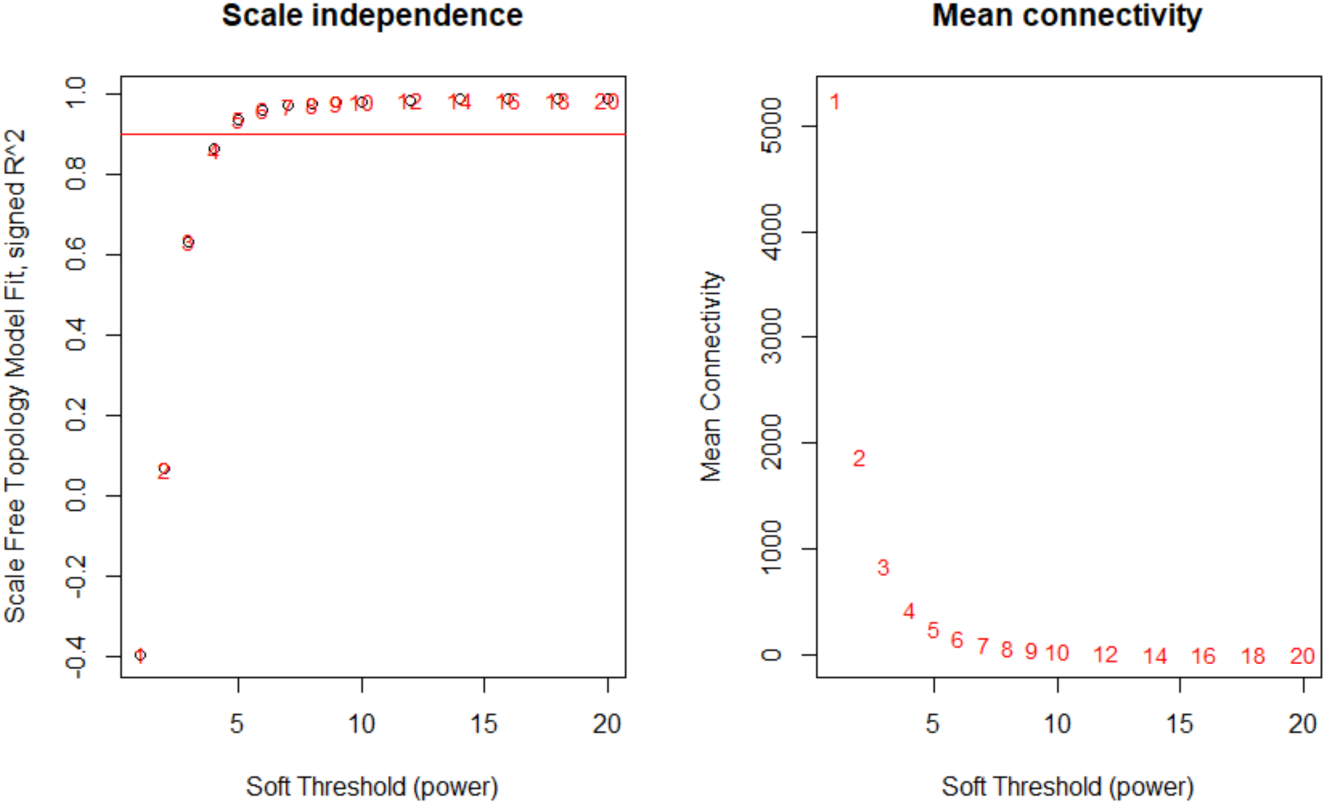
Estimation of Soft Thresholds for model fitting. Left: a scale independence plot depicting the relationship between the soft threshold (x-axis) and the scale-free topology model fit, measured by the signed R² (y-axis); the threshold line, indicated between 4 and 5, assists in identifying an appropriate soft threshold for achieving optimal scale independence. Right: mean connectivity plot illustrating the association between the soft threshold (x-axis) and mean connectivity (y-axis).

**Figure 5.**
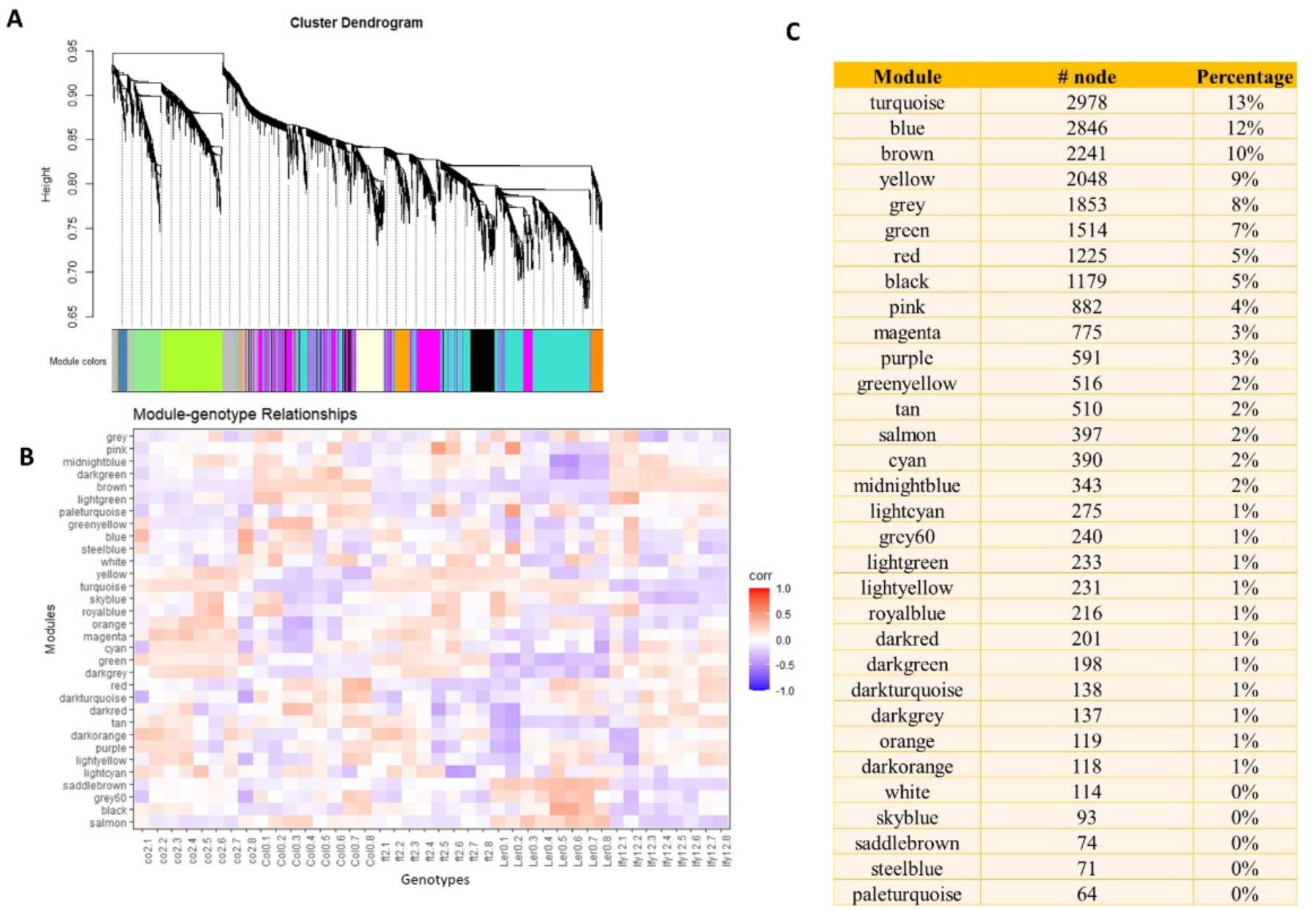
Identification of WGCNA Modules. (A) Hierarchical clustering dendrogram of modules obtained through Weighted Gene Co-expression Network Analysis (WGCNA); each leaf node corresponds to a gene, and branches group genes into distinct modules based on their co-expression patterns; color bands at the bottom of the dendrogram indicate different modules, reflecting sets of genes with similar expression profiles; colors are assigned arbitrarily to aid visual identification. (B) Module-genotype relationship heatmap illustrating the relationships between gene expression modules (rows) and genotypes (columns); color intensity reflects the degree of correlation, with warmer tones indicating stronger positive associations. (C) Modules resolved, with the number of nodes in each and their % contributions to the total.

To relate this modularization to the known genetic regulators of flowering in Arabidopsis, we identified a set of 77 genes previously implicated in the flowering process (Data S1). These genes, which we termed seed or ‘core’ genes, were assigned to our gene modules (Table 2). Interestingly, the 77 genes were distributed across 18 modules, but with 60% located within just five: *Black, Red, Turquoise, Blue*, and *Purple*.

**Table 2.**
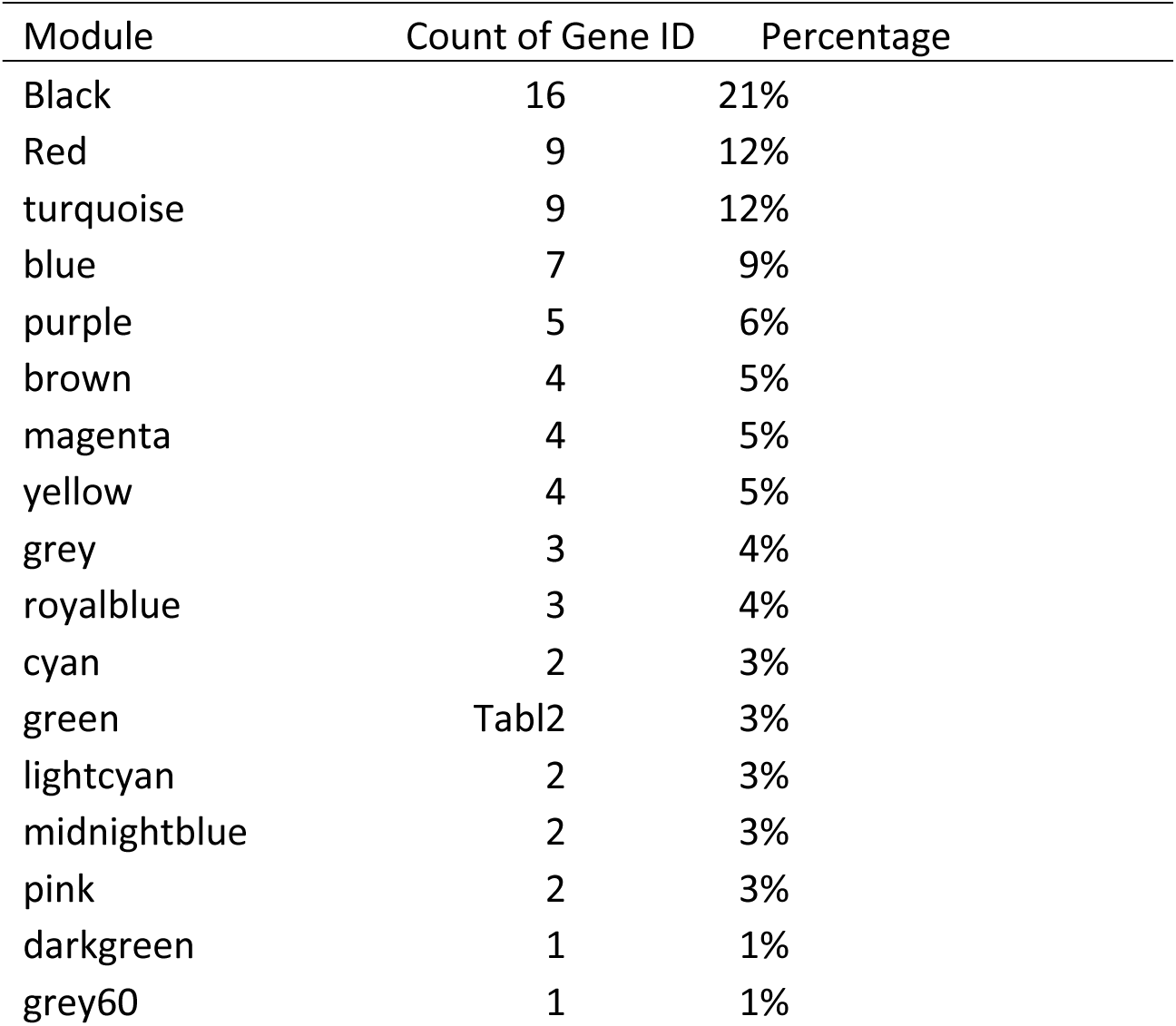
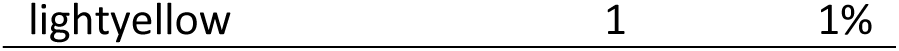
Module attribution of core genes in the flowering time network.

### Ranking of control nodes within flowering subnetworks

It is expected that the most important genes for control of the flowering transition will play diverse roles across pathways at different plant life stages. It is therefore unrealistic to expect that a single algorithm, even one as robust as that employed in WGCNA, will be able to correctly ascribe all regulatory genes to a single module. To address this, we applied a localized adaptation of the Hierarchical Complete Linkage Clustering (HCLC) algorithm to assign the core genes to a central subnetwork, which we termed the “flowering subnetwork” or “flowering module.” We then appended all network nodes which lay within a unitary distance of one of more of these core nodes to the flowering subnetwork, to provide a full representation of how the core nodes can influence the wider molecular landscape via their neighbors. To ensure controllability in measuring controls for the nodes, a directed network comprising 763 nodes and 21,459 edges was constructed. Among these nodes, 326 serve as regulators (Data S1). The degree distribution, a crucial indicator of node connectivity, was found to follow a scale-free distribution (Figure 6). To identify those nodes that exert the greatest control over flowering processes, we used the 326 identified regulator genes to assess the network’s controllability by calculating their individual controllabilities, and the centrality metrics of their associated nodes (Table 3).

**Figure 6.**
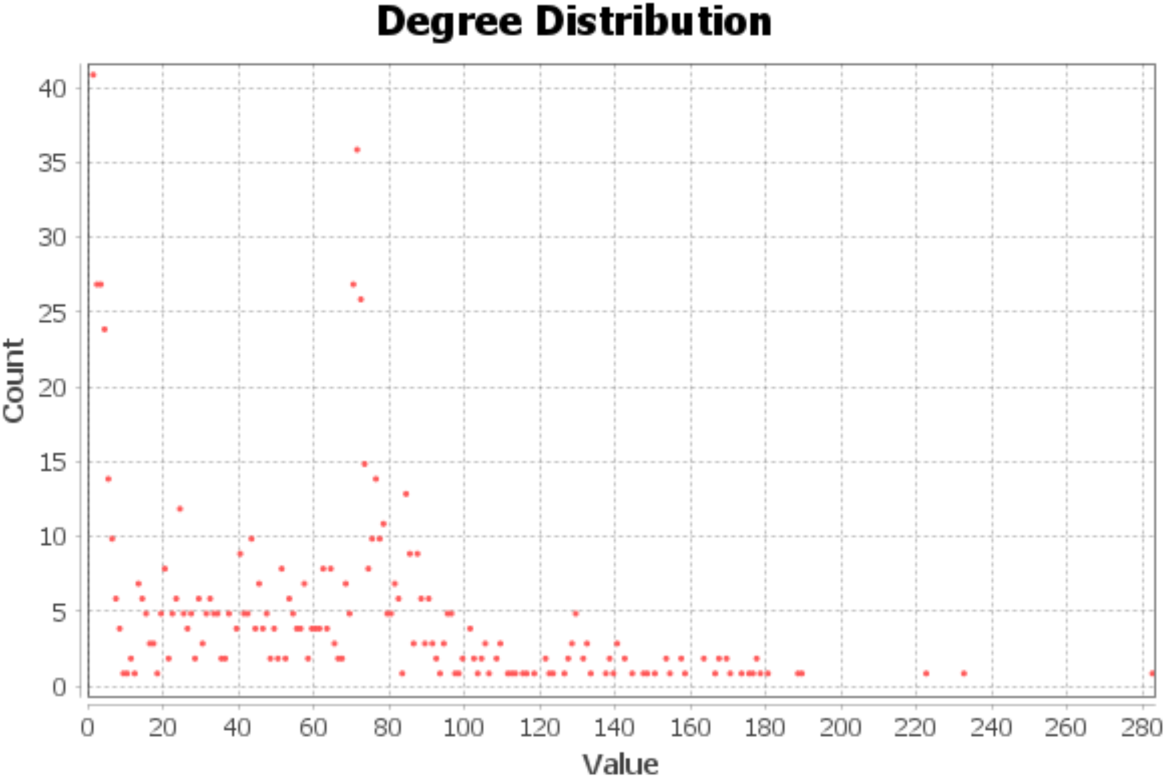
Degree Distribution Plot for the Arabidopsis flowering time network. The x-axis represents the degree values (number of connections) for each node, and the y-axis the corresponding frequency or count of nodes with a given degree.

**Table 3.** Characteristics of the reconstructed regulatory network for flowering.

| Measure | Result |
| --- | --- |
| Nodes | 763 |
| Edges | 21459 |
| Average Degree | 28.125 |
| Network Diameter | 18 |
| Graph Density | 0.037 |
| Average Path Length | 2.741 |

As expected, this approach revealed a hierarchy of gene influence within the network. Notably, a single gene was resolved as the most controllable, overseeing the regulation of 10 nodes within the network. Following closely, 13 genes each exhibited the capability to control nine additional nodes, while another set of 17 genes demonstrate control over eight nodes each (with very large numbers controlling smaller subsets). We conclude that 31 genes within this regulatory network possess the potential to exert significant control, collectively governing a substantial portion of the flowering network, encompassing the remaining 263 genes.

### Driver genes of flowering form a control network

Having identified sets of 77 core genes and 31 controller (or driver) genes, we further considered the functional implications of these gene sets, their associated pathways and biological functions. We determined the closeness centrality scores for each gene as proxies for the strength of regulatory control they exert over the network (Data S2) and ranked them. The most influential genes were resolved as AT2G03710, AT5G10140, and AT3G54340, with degrees of 38, 34, and 30, respectively, while AT2G03710, AT3G54340, and AT1G01060 exhibited the highest closeness centrality scores, measuring 0.57513, 0.563452, and 0.555, respectively, indicating high overlap between these approaches (Table 4).

**Table 4.** Driver genes in the flowering regulatory network, and their respective controllability powers.

| Network Characteristics | High Network Coverage | High Controllability |
| --- | --- | --- |
| AT1G01060 | AT2G03710 | AT3G10480 |
| AT3G54340 | AT1G19350 | AT1G27360 |
| AT5G10140 | AT3G45150 | AT1G19180 |

We further computed various network metrics, including harmonic closeness centrality, betweenness centrality, bridging coefficient, bridging centrality, Authority, Hub, PageRank, and the count of triangles for this regulatory network (Data S2). The amalgamation of these metrics pinpointed nine genes within the network that play a pivotal role in the flowering process (Table 4). An examination of the topological structure of this network section revealed the interplay among these factors which can be considered to form a putative ‘Transition Control Unit (TCU)’ (Figure 7): strikingly, this TCU integrates the thermosensory, light, and internal pathways (including the autonomous pathway [24]) and the circadian clock [49]. We conclude that the TCU encompasses the smallest possible number of genes that are necessary to form a common regulatory network for these flowering pathways.

**Figure 7.**
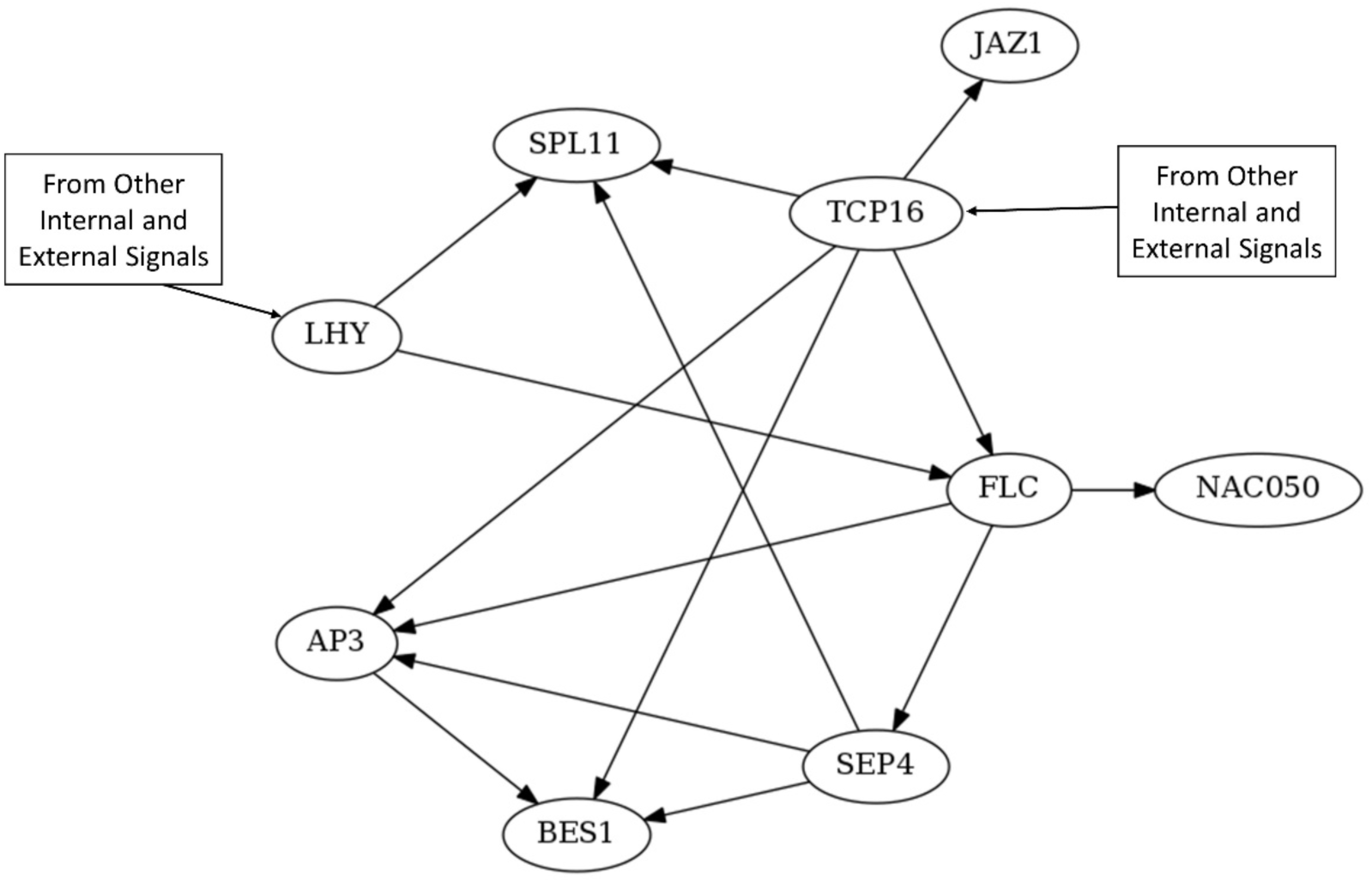
The transition control unit (TCU) for vegetative to reproductive transition in *Arabidopsis thaliana*. Arrows indicate regulatory relations between genes and hence flow of information through the network.

**Table 4.** Key nodes in the regulatory network, identified based on network characteristics (Coverage and Controllability).

| Driver Node | Number of controlled Nodes |
| --- | --- |
| AT3G10480 | 10 |
| AT1G18570, AT1G19180, AT1G19485, AT1G27360, AT1G30210, AT4G00050, AT5G08330, AT5G20570, AT5G36657, AT5G44160, AT5G49400, AT5G56330, AT5G60200 | 9 |
| AT1G10470, AT1G17280, AT1G20140, AT1G48610, AT1G72300, AT2G01930, AT2G21320, AT2G38090, AT2G44745, AT2G46970, AT3G04730, AT3G12250, AT3G50410, AT4G25440, AT5G05190, AT5G08070, AT5G25930 | 8 |

### Features of flowering control by *LHY* and *TCP16*

Visual examination of the nine-gene TCU suggests that two genes, *LHY* and *TCP16*, are positioned at the points at which information from the external flowering time pathways enters the network (Figure 7). To test the implication that these may be key integrators of the flowering transition, we determined the expression patterns of both genes across Arabidopsis growth stages 3.5 to 6, corresponding to the period from which 50% of adult rosette leaves reached full growth, to the initiation of floral bud appearance (Table 1). Given that transcription levels of *LHY* follow a well-established diurnal oscillation connected to its roles in the circadian clock (e.g. [50–51]), levels were monitored across the day-night cycle. Three temperature levels (15, 23, and 28 °C) were applied in the experimental design. As anticipated, plant growth was retarded at lower temperatures, accompanied by a noticeable elongation of each growth stage (Figure 8): plants subjected to cold treatment took 8.4 days longer to reach growth stage 6, while those under heat treatment reached it 2.3 days earlier (Table 5). Importantly, we normalized our data against growth stage rather than chronological day. We found that the average expression of both genes tended to increase across these growth stages (Figure 8) and oscillated across the day-night cycle (Figure 8B). As noted above, this was expected for *LHY* but had not been reported for *TCP16. TCP16* had however been predicted to show diurnal oscillations by Andrés-Colás et al. who found that its target *COPT3* oscillated, while *TCP16* levels were elevated at dawn [52]. Our data confirms the oscillatory expression of *TCP16* transcripts and shows that they do indeed peak at dawn, in fact when *LHY* expression is at its lowest (Figure 8). Examining the general trend of the transcriptional levels depicted in Figures 8A-C, it is noticeable that the expression of *LHY* at any given time in the diurnal cycle initially increases through developmental stages, decreases, then shows another, more modest increase; this pattern is consistent across treatments. However, the area under the curves differs, as do their mean gradients at the turning points (Table 5): this indicates that the pace of expression changes differs across in each day for each treatment. This complexity cannot be explained by simple linear regression models (Figure 8D-F). Together with the expression behaviour of *TCP16* (Figure 8), we conclude that the two genes act together to form a ‘double oscillator’ at the heart of the TCU of flowering time network.

**Figure 8.**
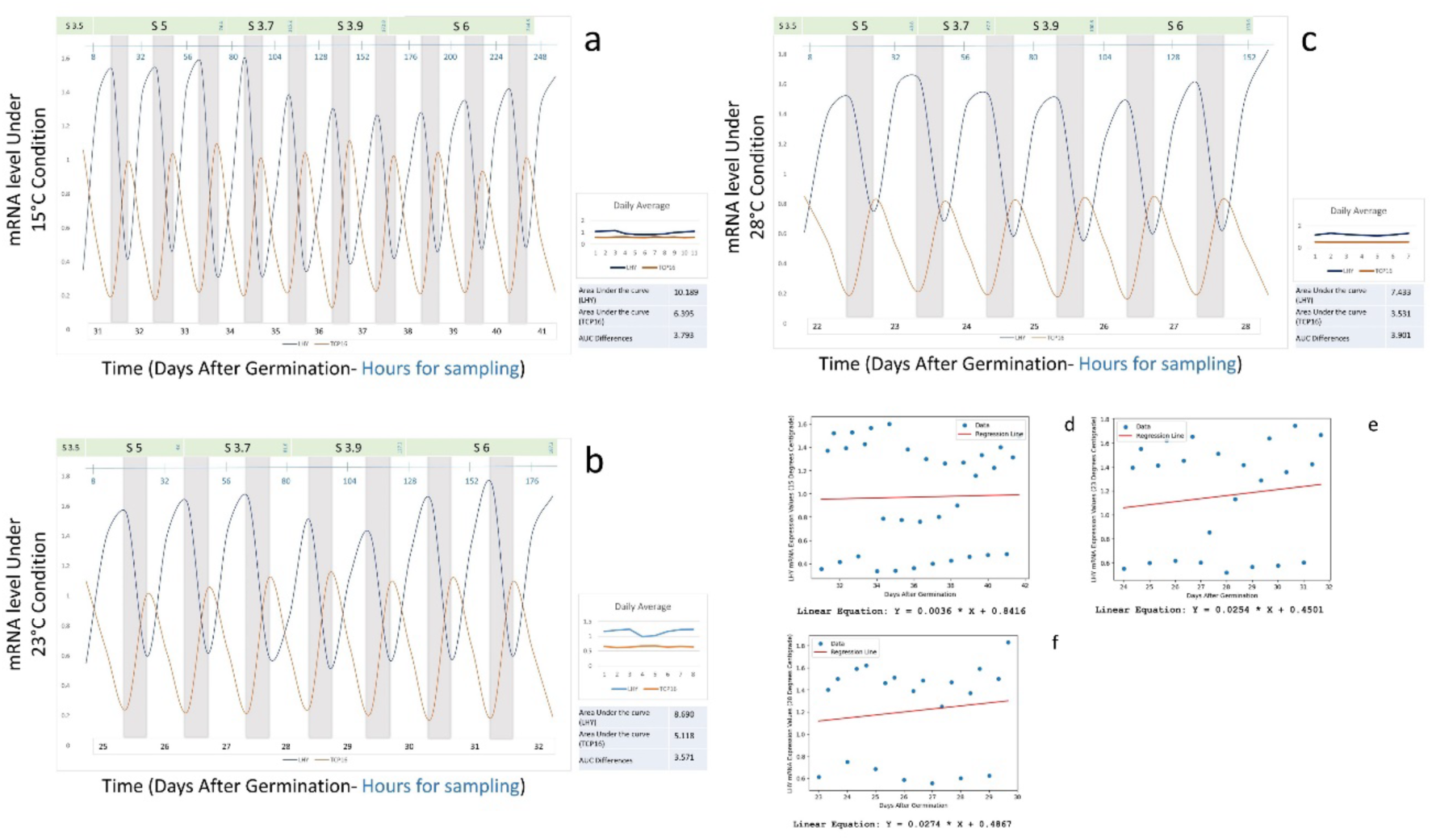
Transcriptional behavior of LHY and TCP16, the two key members of the Flower Transition Gene Regulatory Network (FT-GRN) in Arabidopsis within the transition control unit (TCU). Expression patterns are depicted in blue (*LHY*) and orange (*TCP16*) and are delineated from stage S5 to S6 across diurnal cycles and under various temperature treatments: 15 °C (a, left), 23°C (b, left), and 28 °C (c-left). The average daily expression profiles are shown in composite graphs (a-c, right), with the area under each expression curve shown as a quantitative measure of daily expression changes. (d-f), linear models of the responses of *LHY* across temperatures (15°C, 23°C, and 28°C, respectively).

**Table 5.** The differences in the duration of each growth stage in different temperature treatments; * indicates the number of days from germination to the end of Stage 3.5.

|  |  | 15°C | 23°C | 28°C |
| --- | --- | --- | --- | --- |
| Duration (Days) | *S3.5 | 30 | 24 | 22 |
|  | S5 | 3.1 | 2 | 1.7 |
|  | S3.7 | 1.7 | 1.4 | 1.1 |
|  | S3.9 | 2.4 | 1.9 | 1.4 |
|  | S6 | 3 | 2.5 | 2.2 |
|  | Sum | 40.2 | 31.8 | 28.4 |
| Area Under the curve |  | 10.18935047 | 8.69033335 | 7.43366504 |
| Average Slope of the curve<br>at the turning points |  | -0.04016 | 0.093215 | 0.017504 |

## Discussion

In this study, we have used the extensive knowledge of the molecular biology controlling the flowering transition in Arabidopsis to model the interactions between flowering time genes and have identified a central control module which integrates information from many endogenous and exogenous pathways. Within this central control module, we identified *LHY* and *TCP16* as key to this integrative function. *LHY* has many well-known roles in flowering and development transitions and might have been predicted *a priori* as a likely integrator of flowering cues, while *TCP16* is a much more obscure gene. Our modelling approach has therefore validated previously predictions while identifying potentially fruitful avenues for future research. Using this knowledge to re-engineer flowering time control could bring major benefits for increasing output of new varieties by seed production companies and optimizing seed storage processes [53] and make cropping systems more robust to climate challenges by matching flowering time to optimal times for avoiding environmental stresses while increasing their diversity by synchronizing the flowering of plants from different geographies [54].

The experiment clearly demonstrates the significant impact of genotype, temperature, and light intensity on the flowering time of *Arabidopsis thaliana*. Wild-type genotypes (Col-0 and Ler) exhibited expected trends where higher temperatures and light intensities promoted earlier flowering. This aligns with previous findings that temperature and light intensity are critical environmental factors regulating flowering time [55–56].

### Development of a Position Weight Matrix-based whole-transcriptome model of the flowering transition network

Arabidopsis flowering time genes have been studied from multiple perspectives, and many genes involved in flowering time have been identified over the last few decades (see e.g. [57–59]. It has become increasingly clear that the pathways in which these genes act interact with each other, and with the wider environment, in complex ways: for example, connections have been identified between flowering genes and the thermosensory [56] and light-responsive [60] pathways, while proteins such as CONSTANS form regulatory links with the circadian clock [61–62]. However, understanding how the information which these genes and pathways transmit is integrated to control the plant’s vegetative to reproductive shift remains challenging (Figures 1-2), leading to a growing focus on network modelling approaches [ 8, 43, 63]. For example, Chávez-Hernández et al. (2022) determined the phase transition within the SAM as a key regulator in the flowering transition [10]. In this study, we extended studies such as these by making use of genome-wide transcriptional data during the flowering transition, derived from different genotypes under different conditions to allow dynamic, whole-genome networks to be modelled (Figures 3-4).

In contrast to the use of the Flowering Transition Gene Regulatory Network (FTGRN) method used by Chávez-Hernández and colleagues, our Position Weight Matrix (PWM)-based method is less rooted in literature-based network construction and Boolean modeling. While FTGRN has been instrumental in uncovering qualitative dynamics within the flowering transition gene regulatory network, our PWM-based method extends this to by enhancing the nature of the genetic connections model, incorporating position weight matrices derived from sources such as CisBP and JASPAR to ensure comprehensive coverage of TF motifs. Furthermore, our method incorporates tests of statistical significance, via enrichment analysis and corrections for multiple testing. The PWM-based method’s binary representation of binding sites, while a potential limitation in capturing quantitative dynamics, also offers a clear advantage in its ability to decipher specific nucleotide preferences: this precision is critical for understanding the dynamics of TF binding and subsequent impact on gene expression, which may be less detailed in Boolean models. Since our dataset took the form of a time-course of sufficient sample size, we were also able to derive a co-expression network: by applying a multi-layered strategy (via Pearson correlations) we could not only ensure the inclusion of flowering genes but also indicate levels of hierarchy within the co-expression network.

### The main Arabidopsis flowering pathways converge on a single control module

By applying our approach, all genes responsive to flowering could be arranged into modules (Figure 5), regardless of what flowering time phenotypes, if any, had previously been associated with them. The structure which emerged suggests potential clustering of interacting regulatory genes: the prominence of the *turquoise* module, for instance (Figure 5B), further exploration of which could identify key elements in flowering processes specific to the Col-0 and L*er*-1 genotypes. An addition source of novelty in our approach was the use of the Gephi program to visualize the identified networks, including their structural properties and the levels of connectivity within its topology. The observation that the flowering module network defined by HCLC follows a scale-free distribution (Figure 6) substantiates the scale-free nature of the flowering regulatory network and likely contributes to the inherent robustness and adaptability of the system.

After determining the flowering time information system (*sensu* Tkačik and Walczak [64]), we identified the most important nodes for network controllability via centrality measures. This allowed us to identify the nodes in which the information from different paths enters them (the entry channels/nodes) and passes to the controller nodes in the transition system boundary (as shown schematically in Figure 2). In this way, various environmental signals may affect the network from dozens of different paths, and these internal and external signals are processed in various processing units and directed to other units including what we termed the [vegetative-reproductive] transition control unit (TCU) through one or more channels to be translated to the physiological response. This simultaneous assessment of both gene centrality and influence, in the context of a spectrum of network metrics, strengthens our identification of key regulatory genes within the dynamics of flowering time regulation (Tables 4-5). Notably, all the main flowering pathways converge in this central TCU: while it was previously demonstrated that the autonomous and thermosensory pathways share common regulators [65], our model indicates that all pathways related to flowering are at least partly shared and interconnected in this network unit. This makes sense, given the need for the plant to respond to the totality of its environment in order to enter flowering at the evolutionarily optimal point. Examination of Figure 7 further identifies two bottlenecks in these regulatory units, the genes *LHY* and *TCP16,* through which the information inputted from the environmental and internal pathways is further canalized, before being distributed among seven other genes (*JAZ1*, *SPL11*, *AP3*, *FLC*, *SEP4*, *BES1* and *NAC050*). The nature of the interactions which underlie these layers of genetic control – none of these necessarily involve physical interaction, or even co-expression within the same cells (and indeed the control of *FLC* by *LHY* is known to act indirectly via *VIN3* [66] – represent interesting directions for future study.

### Novel roles for *TCP16* and *LHY* as a potential Double Oscillator system integrating information during the flowering transition

The key positions of *LHY* and *TCP16* in the topology of the Transition Control Unit (TCU) suggests that their transcriptional activity may control the outputs of the flowering pathways. It should be noted that the existence of nine genes in this submodule does not mean that only these nine genes are responsible for regulating flowering time. Rather, there are more connections between these genes and other genes involved in the process of transition from the vegetative to reproductive stage, and this network represents the connection between the most important nodes in the regulatory network. The most important nodes are the nodes that have received a high score in terms of controllability and centrality. For example, LHY and CCA1 genes have long been known as two important members of the circadian clock [49], but no trace of CCA1 can be seen in this TCU. The reason for this is not the absence of this gene in the network, but the reason is the lack of identifying it “alone” to exert control over the network or its less centrality in the regulatory network.

The TCP16 transcription factor, a member of the Class I TCP family in Arabidopsis thaliana [67–68], is implicated in the regulation of diverse developmental processes, including the timing of flowering ([69–70] and references therein). Although the precise downstream targets of TCP16 involved in flowering time control have yet to be fully characterized, evidence from related TCP family members suggests several potential pathways through which TCP16 may exert its influence in repression meristem identity [71] and copper transport, with particularly important roles in the pollen [52].

One potential target of TCP16 is the *FLOWERING LOCUS T* (*FT*) gene, a critical component in the photoperiodic regulation of flowering. *FT* encodes the florigen, a mobile protein signal that promotes flowering by inducing the expression of key floral identity genes in the shoot apical meristem. TCP transcription factors, including TCP18 (BRANCHED1), have been shown to repress FT expression, delaying flowering [72]. Given the functional similarities within the TCP family, it is plausible that TCP16 may similarly modulate FT expression, thereby influencing the timing of flowering. TCP16 may also interact with components of the circadian clock, a key regulator of flowering time. The circadian clock controls the expression of *CONSTANS*, which in turn regulates *FT* [4]. TCP proteins are known to integrate environmental signals, such as light, into developmental processes [73], suggesting that TCP16 could modulate flowering time by influencing circadian clock-regulated pathways. Additionally, TCP16 might regulate GROWTH-REGULATING FACTORs (GRFs), which are transcription factors involved in various developmental processes, including leaf development and flowering. GRFs can affect the expression of genes involved in flowering [74], indicating that TCP16’s regulation of GRFs could be another mechanism through which it controls flowering time.

Hormone signalling pathways, particularly the gibberellin (GA) pathway, represent another potential avenue for TCP16’s role in flowering time control. Gibberellins promote flowering, especially under non-inductive photoperiods [75]. TCP16 may influence the expression of genes involved in GA biosynthesis or signalling, thus integrating hormonal cues into the regulation of flowering. Finally, TCP16 may interact with the age pathway through the regulation of *SQUAMOSA PROMOTER BINDING PROTEIN-LIKE* (*SPL*) genes. *SPL* genes are regulated by the microRNA miR156 and promote flowering by activating *FT* and other flowering-related genes [76]. Given the role of TCP proteins in developmental timing [77], TCP16 might influence flowering time by modulating SPL gene expression or miR156 activity.

In contrast, *LHY* encodes a core component of the circadian clock [78] which is already known to control flowering in Arabidopsis [46, 49, 79]. Mutation of *LHY* leads to organismal arrhythmia and late day-length-insensitive flowering [56], repressing flowering under normal light conditions but accelerating it under continuous light [49, 80]. *LHY* has also been shown to interact with the vernalization pathway [66], playing roles in the layers of crosstalk known to exist between circadian regulation and environmental signalling [51]. In *Ziziphus jujuba,* both endogenous and ectopically-expressed Arabidopsis *LHY* were supressed by ZjSEP3, leading to accelerated flowering which could be rescued by expression of *LHY* under the 35S promoter [81].

LATE ELONGATED HYPOCOTYL (LHY), a key component of the circadian clock in *Arabidopsis thaliana*, plays a critical role in regulating various physiological processes, including flowering time. LHY functions as a transcription factor that integrates circadian rhythms with developmental and environmental signals. Understanding its downstream targets provides insights into how circadian regulation intersects with flowering time control. One primary target of LHY is the CONSTANS (CO) gene, which is crucial for photoperiodic flowering. CO encodes a transcription factor that activates the FLOWERING LOCUS T (FT) gene, a major signal for floral induction [82]. LHY directly represses CO expression in the evening through its role in the circadian clock, thereby modulating the timing of flowering in response to day length [83]. The rhythmic expression of LHY and its control over CO exemplify the integration of the circadian clock with flowering time regulation. FT itself is another key downstream target regulated by LHY. FT acts as a mobile signal that promotes flowering by triggering the expression of floral meristem identity genes [84]. LHY’s regulation of CO, which in turn affects FT expression, underscores its central role in coordinating flowering time with circadian rhythms. LHY also interacts with genes involved in the circadian clock machinery, such as *CCA1* (*CIRCADIAN CLOCK ASSOCIATED 1*) and *PRR* (*PSEUDO-RESPONSE REGULATOR*). *CCA1* and genes of the *PRR* family, including *PRR7* and *PRR9*, are integral to the circadian feedback loop, influencing the expression of downstream targets like *CO* and *FT* [78]. By regulating these clock genes, LHY modulates the overall rhythmic output of the clock, which has downstream effects on flowering time.

The influence of LHY on hormone signaling pathways further illustrates its role in flowering time control. Gibberellins (GAs), which promote flowering under non-inductive photoperiods, are modulated by LHY through its regulation of hormone-responsive genes [85]. For example, LHY affects the expression of genes involved in GA biosynthesis and signaling, linking circadian control with hormonal regulation of flowering. *SPL* genes are also affected by LHY, as well as by the microRNA miR156, which is itself under circadian control [86]. By affecting *SPL* expression through its influence on miR156, LHY impacts the transition from vegetative to reproductive growth. In summary, LHY regulates flowering time through a network of downstream targets, including CO, FT, circadian clock components, and hormone signaling pathways. Its role in integrating circadian rhythms with developmental and environmental signals highlights the complexity of flowering time regulation. Further studies using genome-wide approaches, such as ChIP-seq and transcriptomic analysis, will provide a deeper understanding of the specific genes and pathways regulated by LHY, elucidating its precise role in flowering time control.

Our analysis of the transcriptional responses of these genes indicates the core regulators of the flowering time TCU is characterized by oscillations of both genes (Figure 8). Potentially, this could imply the existence of a double oscillation system [87–90] in which *LHY* and *TCP16* reciprocally influence is affected by upstream signals from internal and external signals modulate these signals and respond to them by changing the expression behaviour in a cyclical manner, affecting downstream elements either through direct TF activity or other, more remote effects (including non-cell autonomously). Although double oscillations are observed within the clock for LHY/BES1/TPL-CCA1 [91], our model does not however predict *LHY* and *TCP16* to affect each other directly (Figure 7): their behaviour is instead connected through other information flow paths. This agrees with the finding of Leal Valentim and colleagues that flowering genes can have their maximal effect on genes which they interact with only indirectly [8].

Finally, Figures 8 reveals that plants at different growth stages reach the peaks and troughs of gene expression at different times under different environments, along with other distinct characteristics among the waves. This reinforces that gene expression studies based on hourly or daily sampling (rather than growth stage-based sampling) may lead to misleading results. For instance, comparing the average expression of *LHY* in two different treatments (Figure 8) suggests an increase in gene expression upon reaching stage S6. However, relying on linear trends could result in erroneous conclusions, as the 15 °C treatment delays progression to stage S6. Hence, the peak of the fourth wave observed in the cold treatment is several days behind the equivalent under 23 °C conditions (Table 1, Figure 8). It might mistakenly be inferred that the expression level of *LHY* is higher in the 15 °C treatment than in the 23 °C treatment. Even in a specific stage like S6, comparing gene expression between the two treatments is challenging, as this comparison essentially reveals points of increase or decrease in expression. However, in practice, factors such as the amplitude of the wave and the area under the waveforms also differ (as calculated according to the Riemann method; Methods). This suggests that other complex mechanisms are involved in regulating gene values at each time, potentially including miRNAs and mRNA stability [74, 92], all of which should be considered for testing the impact of this genetic network at the tissue and organismal level. Ultimately, our data suggests that if oscillation management circuits in *LHY* and *TCP16* could be designed and implemented, it would be possible to engineer flowering time in Arabidopsis.

## Methods and Materials

### Data acquisition and handling

*Expression data.* Transcriptomic data from 40 samples of five genotypes of *Arabidopsis thaliana* were accessed from GEO database Series GSE576. These were derived from wild-type Columbia (Col-0) and Landsberg *erecta* (L*er*), and the mutants *leafy*-*12* (*lfy-12*, in the Col-0 background), *constans-*2 (*co-2*, in the L*er* background) and *flowering locus T-2* (*ft-2*, also in L*er*). In each case, data was derived from microarrays monitoring gene expression of 22,810 genes and were performed as a time-course after the plants had been moved from short day to long day growth conditions (0, 3, 5, and 7 days after the shift); each genotype is represented by two replicates.

*Weighted Gene Co-expression Network Analysis*. Gene expression data was checked for quality and used to construct a gene co-expression network in the Weighted Gene Co-expression Network Analysis (WGCNA) R package, where the pairwise gene similarity matrix was calculated based on expression profiles. The similarity matrix was transformed into an adjacency matrix using the soft-thresholding method within the package, capturing the weighted nature of gene co-expression relationships. A topological overlap matrix was calculated, hierarchical clustering performed, and co-expression modules identified. The resulting modules were represented by distinct colors.

*Development of the Co-Expression Module*. To construct a co-expression module specifically linked to flowering processes, we employed a refined approach by adapting the Hierarchical Complete Linkage Clustering (HCLC) method, with Pearson correlation serving as the primary metric for assessing gene expression relationships. Regulatory relationships in the constructed network were predicted following Kulkarni et al. [93]. Briefly, position weight matrices (PWMs) representing 916 transcription factors (TFs) were derived from diverse sources, including CisBP, Franco-Zorrilla et al., Plant Cistrome Database, JASPAR 2016, UNIPROBE, AGRIS, and AthaMAP. These PWMs encompass 1793 Arabidopsis motifs, either in PWM or consensus sequence formats, converted into position count matrices scaled to 100. TFs are assigned to gene families based on the PlnTFDB 3.0 database. Gene regions, including long, intermediate, and short segments, were extracted relative to the translation start and end sites. PWMs were then mapped to these gene regions using Cluster-Buster, retaining family-specific information within the binding sites (BS). Enrichment statistics were calculated using the hypergeometric distribution, evaluating motif enrichment in input gene sets. The q-value of enrichment was determined with the Benjamini–Hochberg correction for multiple hypotheses testing. To assess false discovery rates, control datasets were generated, and a default q-value of 0.05 was set.

### Phase Transition Unit Boundary Detection

*Controllability Analysis*. To calculate the centrality of control for nodes, we used an improved version of the method we previously developed [94]. To calculate the controllability of nodes, we modelled the network dynamics through a linear differential equation with a connectivity matrix A, where the vector x(t) represents the state of N nodes at time t. The driver nodes, represented by the matrix B, were identified, and the controllability of the network is determined from the Kalman condition. We then determined the number of nodes which each node controls by considering path lengths between pairs of nodes.: nodes with the highest control centrality (HCC) were identified, constituting the initial set of nodes employing a free-rider strategy. Performance assessment involved comparing these nodes with three other groups: nodes with the highest out-degree, locally selected nodes, and randomly selected nodes. The method incorporates a public goods game with a linear benefit function, with a sigmoid function used to calculate the benefit function, capturing the non-linear behavior often observed in biological systems. Both deterministic and stochastic update rules were applied to model the evolution of strategies in the public goods game, adding a level of uncertainty controlled by the parameter β.

*Architectural Studies*. The Gephi software package (https://gephi.org/; [95]) was employed for the architectural analysis of the constructed networks including node properties, such as degree, betweenness centrality, and clustering coefficients, which were visually encoded to discern key nodes and network modules.

### Validation of Gene Expression

*Plant Materials*. Col-0 and Ler accessions (kindly provided by Dr. Adriana Garay Arroyo from Lab. de Genética Molecular, Epigenética, Desarrollo y Evolución de Plantas. Instituto de Ecología, UNAM), were grown under controlled conditions with light at 100% (120-150 µmol.m^-^².S^-1^) or 60% (70-90 µmol.m^-^².S^-1^) intensity and 15, 23, and 28 °C for day temperature and 13, 21 and 26 °C for night. Plants were grown initially in short days (9 hours light, 15 hours dark) and then transferred to long days (16 hours light, 8 hours dark). To synchronize our sampling, the developmental stages of plant growth were monitored from stages 3.5 to 6.1 (Supplementary File 2). Samples were collected every 6 hours, and bud samples (13-15 per sample) were also taken to assess gene expression levels in the Col-0 genotype. The experimental design employed a completely randomized block design, with two replicates considered for each.

*Quantitative Real Time PCR*. Total RNA was isolated according to Oñate-Sánchez and Vicente-Carbajosa [96]. RNA concentrations were determined on a NanoDrop spectrophotometer (WPA Biowave II+); only samples exhibiting a 260:280 ratio between 1.8 and 2.2 were analyzed further. RNA integrity was determined via agarose gel electrophoresis (0.8%) in tris/borate/ethylene diamine tetra acetic acid (TBE) buffer. 1 μg of total RNA from each sample was used for reverse transcription with the MMLV-RT first-strand cDNA synthesis kit (Thermo Fisher Scientific, USA), used according to manufacturer’s instructions. Quantitative Real-Time PCR was performed on a BioRad instrument (BioRad, USA) with Eva Green Real-Time PCR master mix (Solis BioDyne, Estonia); primers were designed using Oligo 7 (Molecular Biology Insights, USA; Table S1). qRT-PCR cycles consisted of 95 °C for 15 min and 40 cycles at 95 °C for 15 s, 60 °C for 20 s, and 72 °C for 20 s. Relative expression levels were double-normalized against the Ct value of *GLYCERALDEHYDE-3-PHOSPHATE DEHYDROGENASE* (*GAPDH*) as the internal control. qRT-PCR data was analyzed with BioRad CFX Manager Software Ver. 1.6 (BioRad, USA), using the formula (1 + E)^-ΔΔCt^.

**Supplementary Table S1.**
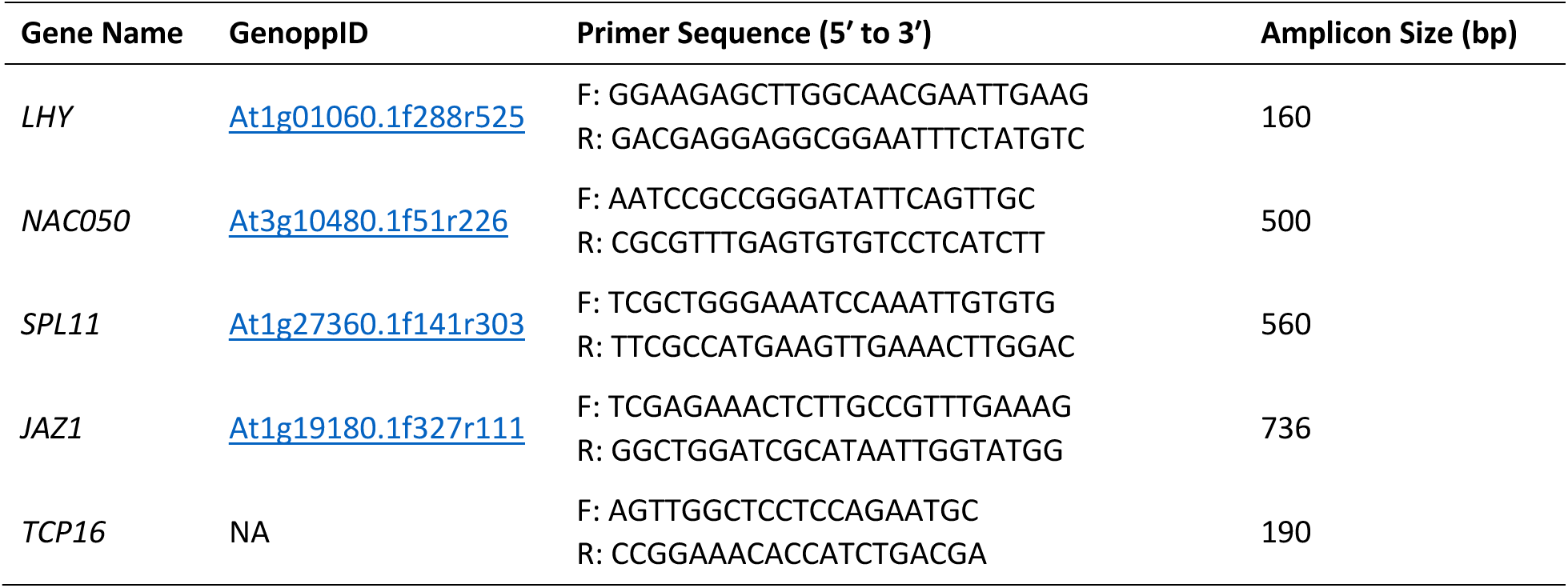
Primer sequences used for gene amplification by qRT-PCR. Primer design. The amplification efficiency of each primer pair was calculated with cDNA serial dilutions using E = 10^-1^/slope-1; only those primer pairs with an efficiency range of 90-110% were selected.

## References

[1] Cho LH, Yoon J, An G. (2017) The control of flowering time by environmental factors. Plant J. 90(4):708–719. doi: 10.1111/tpj.13461.

[2] Yan, W., Wang, B., Chan, E., & Mitchell-Olds, T. (2021) Genetic architecture and adaptation of flowering time among environments. New Phytologist, 230(3), 1214–1227.

[3] Maeda AE, Nakamichi N. (2002) Plant clock modifications for adapting flowering time to local environments. Plant Physiol. 190(2):952–967. doi: 10.1093/plphys/kiac107.

[4] Andrés, F., & Coupland, G. (2012). The genetic basis of flowering responses to seasonal cues. Nature Reviews Genetics, 13(9), 627–639.

[5] Khodorova, N. V., & Boitel-Conti, M. (2013) The role of temperature in the growth and flowering of geophytes. Plants, 2(4), 699–711.

[6] Hwang, K., Susila, H., Nasim, Z., Jung, J. Y., & Ahn, J. H. (2019) Arabidopsis ABF3 and ABF4 transcription factors act with the NF-YC complex to regulate *SOC1* expression and mediate drought-accelerated flowering. Molecular Plant, 12(4), 489–505.

[7] Wang X, Jiang B, Gu L, Chen Y, Mora M, Zhu M, Noory E, Wang Q, Lin C. (2021) A photoregulatory mechanism of the circadian clock in Arabidopsis. Nat Plants. 7(10):1397–1408. doi: 10.1038/s41477-021-01002-z. Epub 2021 Oct 14. Erratum in: Nat Plants. 2021

[8] Leal Valentim, F., Mourik, S.V., Posé, D., Kim, M.C., Schmid, M., van Ham, R.C., Busscher, M., Sanchez-Perez, G.F., Molenaar, J., Angenent, G.C. and Immink, R.G. (2015) A quantitative and dynamic model of the Arabidopsis flowering time gene regulatory network. PloS ONE, 10(2), p.e0116973.

[9] Bouché, F., Lobet, G., Tocquin, P., & Périlleux, C. (2016). FLOR-ID: an interactive database of flowering-time gene networks in Arabidopsis thaliana. Nucleic Acids Research, 44(D1), D1167–D1171.

[10] Chávez-Hernández, E. C., Quiroz, S., García-Ponce, B., & Álvarez-Buylla, E. R. (2022). The flowering transition pathways converge into a complex gene regulatory network that underlies the phase changes of the shoot apical meristem in *Arabidopsis thaliana*. Frontiers in Plant Science, 13.

[11] Quiroz, S., Yustis, J.C., Chávez-Hernández, E.C., Martínez, T., Sanchez, M.D.L.P., Garay-Arroyo, A., Álvarez-Buylla, E.R. and García-Ponce, B. (2021) Beyond the genetic pathways, flowering regulation complexity in Arabidopsis thaliana. Intl J Mol Sci, 22(11), 5716.

[12] Fernández, V., Takahashi, Y., Le Gourrierec, J., & Coupland, G. (2016). Photoperiodic and thermosensory pathways interact through CONSTANS to promote flowering at high temperature under short days. The Plant Journal, 86(5), 426–440.

[13] Wang, H., Pan, J., Li, Y., Lou, D., Hu, Y., & Yu, D. (2016) The DELLA-CONSTANS transcription factor cascade integrates gibberellic acid and photoperiod signaling to regulate flowering. Plant Physiology, 172(1), 479–488.

[14] Li, X., Lai, M., Li, K., Yang, L., Liao, J., Gao, Y., Wang, Y., Gao, C., Shen, W., Luo, M., & Yang, C. (2024) FLZ13 interacts with FLC and ABI5 to negatively regulate flowering time in Arabidopsis. New Phytologist, 241(3), 1334–1347.

[15] Romero, J.M., Serrano-Bueno, G., Camacho-Fernández, C., Vicente, M.H., Ruiz, M.T., Pérez-Castiñeira, J.R., Pérez-Hormaeche, J., Nogueira, F.T. and Valverde, F. (2024) CONSTANS, a HUB for all seasons: How photoperiod pervades plant physiology regulatory circuits. *The Plant Cell*, p.koae090.

[16] Whittaker, C., & Dean, C. (2017) The *FLC* locus: a platform for discoveries in epigenetics and adaptation. Annual Review of Cell and Developmental Biology, 33, 555–575.

[17] Searle, I., He, Y., Turck, F., Vincent, C., Fornara, F., Kröber, S., Amasino, R. A., & Coupland, G. (2006) The transcription factor FLC confers a flowering response to vernalization by repressing meristem competence and systemic signaling in Arabidopsis. Genes & Development, 20(7), 898–912.

[18] Costa, S., & Dean, C. (2019). Storing memories: the distinct phases of Polycomb-mediated silencing of Arabidopsis FLC. Biochemical Society Transactions, 47(4), 1187–1196.

[19] Choi, K., Kim, J., Hwang, H. J., Kim, S., Park, C., Kim, S. Y., & Lee, I. (2011). The FRIGIDA complex activates transcription of *FLC*, a strong flowering repressor in Arabidopsis, by recruiting chromatin modification factors. The Plant Cell, 23(1), 289–303.

[20] Li, Z., Jiang, D., & He, Y. (2018). FRIGIDA establishes a local chromosomal environment for *FLOWERING LOCUS C* mRNA production. Nature Plants, 4(10), 836–846.

[21] Zhu, P., Lister, C., & Dean, C. (2021) Cold-induced Arabidopsis FRIGIDA nuclear condensates for FLC repression. Nature, 599(7886), 657–661.

[22] Shen, L., Zhang, Y., & Sawettalake, N. (2022) A Molecular switch for FLOWERING LOCUS C activation determines flowering time in Arabidopsis. Plant Cell, 34(2).

[23] Cheng, J. Z., Zhou, Y. P., Lv, T. X., Xie, C. P., & Tian, C. E. (2017). Research progress on the autonomous flowering time pathway in Arabidopsis. Physiology and Molecular Biology of Plants, 23, 477–485.

[24] Wu, Z., Fang, X., Zhu, D., & Dean, C. (2020) Autonomous Pathway: *FLOWERING LOCUS C* repression through an antisense-mediated chromatin-silencing mechanism. Plant Physiology, 182(1), 27–37.

[25] Bajczyk, M., Jarmolowski, A., Jozwiak, M., Pacak, A., Pietrykowska, H., Sierocka, I., Swida-Barteczka, A., Szewc, L. and Szweykowska-Kulinska, Z. (2023) Recent insights into plant miRNA biogenesis: multiple layers of miRNA level regulation. Plants, 12(2), 342.

[26] Dong, Q., Hu, B., & Zhang, C. (2022). microRNAs and their roles in plant development. Frontiers in Plant Science, 13, 824240.

[27] Han, Y., Liu, L., Lei, M., Liu, W., Si, H., Ji, Y., … & Zan, Y. (2025). Divergent flowering time responses to increasing temperatures are associated with transcriptome plasticity and epigenetic modification differences at *FLC* promoter region of *Arabidopsis thaliana*. Molecular Ecology, 34(15), e17544.

[28] Liang, Z., Huang, Y., Hao, Y., Song, X., Zhu, T., Liu, C., & Li, C. (2025). The HISTONE ACETYLTRANSFERASE 1 interacts with CONSTANS to promote flowering in Arabidopsis. Journal of Genetics and Genomics.

[29] Mattioli, R., Francioso, A., & Trovato, M. (2022). Proline affects flowering time in Arabidopsis by modulating FLC expression: a clue of epigenetic regulation? Plants, 11(18), 2348.

[30] Nasim, Z., Fahim, M., Hwang, H., Susila, H., Jin, S., Youn, G., & Ahn, J. H. (2021). Nonsense-mediated mRNA decay modulates Arabidopsis flowering time via the SET DOMAIN GROUP 40–FLOWERING LOCUS C module. Journal of Experimental Botany, 72(20), 7049–7066.

[31] Davidson, E., & Levin, M. (2005). Gene regulatory networks. Proceedings of the National Academy of Sciences, 102(14), 4935–4935.

[32] Alvarez-Buylla, E. R., Benítez, M., Dávila, E. B., Chaos, A., Espinosa-Soto, C., & Padilla-Longoria, P. (2007). Gene regulatory network models for plant development. Current Opinion in Plant Biology, 10(1), 83–91.

[33] Jones, D. M., & Vandepoele, K. (2020) Identification and evolution of gene regulatory networks: insights from comparative studies in plants. Current Opinion in Plant Biology, 54, 42–48.

[34] Del Vecchio D, Dy AJ, Qian Y. (2016) Control theory meets synthetic biology. J R Soc Interface. 13(120):20160380. doi: 10.1098/rsif.2016.0380.

[35] Haque, S., Ahmad, J. S., Clark, N. M., Williams, C. M., & Sozzani, R. (2019). Computational prediction of gene regulatory networks in plant growth and development. Current opinion in plant biology, 47, 96–105.

[36] Ko, D. K., & Brandizzi, F. (2020) Network-based approaches for understanding gene regulation and function in plants. The Plant Journal, 104(2), 302–317.

[37] Schwab, J. D., Kühlwein, S. D., Ikonomi, N., Kühl, M., & Kestler, H. A. (2020) Concepts in Boolean network modeling: What do they all mean? Computational and Structural Biotechnology Journal, 18, 571–582.

[38] Timmermann, T., González, B., & Ruz, G. A. (2020). Reconstruction of a gene regulatory network of the induced systemic resistance defense response in Arabidopsis using boolean networks. BMC Bioinformatics, 21(1).

[39] Pavlinova, P., Samsonova, M. G., & Gursky, V. v. (2021) Dynamical Modeling of the Core Gene Network Controlling Transition to Flowering in *Pisum sativum*. Frontiers in Genetics, 12.

[40] Dondelinger, F., Husmeier, D., & Lèbre, S. (2012) Dynamic Bayesian networks in molecular plant science: Inferring gene regulatory networks from multiple gene expression time series. Euphytica, 183(3).

[41] Castro, J. C., Valdés, I., Gonzalez-García, L. N., Danies, G., Cañas, S., Winck, F. V., Ñústez, C. E., Restrepo, S., & Riaño-Pachón, D. M. (2019) Gene regulatory networks on transfer entropy (GRNTE): A novel approach to reconstruct gene regulatory interactions applied to a case study for the plant pathogen *Phytophthora infestans*. Theoretical Biology and Medical Modelling, 16(1).

[42] Ingkasuwan, P., Netrphan, S., Prasitwattanaseree, S., Tanticharoen, M., Bhumiratana, S., Meechai, A., Chaijaruwanich, J., Takahashi, H., & Cheevadhanarak, S. (2012) Inferring transcriptional gene regulation network of starch metabolism in Arabidopsis thaliana leaves using graphical Gaussian model. BMC Systems Biology, 6.

[43] Wang CC, Chang PC, Ng KL, Chang CM, Sheu PC, Tsai JJ. (2014) A model comparison study of the flowering time regulatory network in Arabidopsis. BMC Syst Biol. 8:15. doi: 10.1186/1752-0509-8-15.

[44] Koornneef, M., Alonso-Blanco, C., Blankestijn-de Vries, H., Hanhart, C. J., & Peeters, A. J. M. (1998). Genetic interactions among late-flowering mutants of Arabidopsis. Genetics, 148(2), 885–892. Retrieved from <Go to ISI>://WOS:000072187500031

[45] Weigel, D., Alvarez, J., Smyth, D. R., Yanofsky, M. F., & Meyerowitz, E. M. (1992). Leafy Controls Floral Meristem Identity in Arabidopsis. Cell, 69(5), 843–859. doi:Doi 10.1016/0092-8674(92)90295-N

[46] Suárez-López, P., Wheatley, K., Robson, F., Onouchi, H., Valverde, F., & Coupland, G. (2001) CONSTANS mediates between the circadian clock and the control of flowering in Arabidopsis. Nature, 410(6832), 1116–1120.

[47] Kardailsky, I., Shukla, V. K., Ahn, J. H., Dagenais, N., Christensen, S. K., Nguyen, J. T., – Weigel, D. (1999). Activation tagging of the floral inducer. Science, 286(5446), 1962–1965. doi:DOI 10.1126/science.286.5446.1962

[48] Kobayashi, Y., Kaya, H., Goto, K., Iwabuchi, M., & Araki, T. (1999). A pair of related genes with antagonistic roles in mediating flowering signals. Science, 286(5446), 1960–1962. doi:DOI 10.1126/science.286.5446.1960

[49] Fujiwara, S., Oda, A., Yoshida, R., Niinuma, K., Miyata, K., Tomozoe, Y., Tajima, T., Nakagawa, M., Hayashi, K., Coupland, G., & Mizoguchi, T. (2008) Circadian clock proteins LHY and CCA1 regulate SVP protein accumulation to control flowering in Arabidopsis. The Plant Cell, 20(11), 2960–2971.

[50] Mizoguchi, T., Wheatley, K., Hanzawa, Y., Wright, L., Mizoguchi, M., Song, H.R., Carré, I.A. and Coupland, G. (2002) LHY and CCA1 are partially redundant genes required to maintain circadian rhythms in Arabidopsis. Developmental Cell, 2(5), 629–641.

[51] Paajanen, P., de Barros Dantas, L. L., & Dodd, A. N. (2021) Layers of crosstalk between circadian regulation and environmental signalling in plants. Current Biology, 31(8), R399–R413.

[52] Andrés-Colás, N., Carrió-Seguí, A., Abdel-Ghany, S. E., Pilon, M., & Peñarrubia, L. (2018). Expression of the intracellular COPT3-mediated Cu transport is temporally regulated by the TCP16 transcription factor. Frontiers in Plant Science, 9, 910.

[53] Kehrberger, S., Holzschuh, A. (2019) How does timing of flowering affect competition for pollinators, flower visitation and seed set in an early spring grassland plant?. Sci Rep 9, 15593

[54] Sajid SS, Hu G. (2022) Optimizing crop planting schedule considering planting window and storage capacity. Front Plant Sci. 13:762446. doi: 10.3389/fpls.2022.762446.

[55] Johanson, U., West, J., Lister, C., Michaels, S., Amasino, R., & Dean, C. (2000). Molecular analysis of FRIGIDA, a major determinant of natural variation in Arabidopsis flowering time. Science, 290(5490), 344–347.

[56] Blažquez, M. A., Green, R., Nilsson, O., Sussman, M. R., & Weigel, D. (1998). Gibberellins promote flowering of Arabidopsis by activating the LEAFY promoter. The Plant Cell, 10(5), 791–800.

[57] Johansson M, Staiger D. (2015) Time to flower: interplay between photoperiod and the circadian clock. Journal of Experimental Botany, 66(3):719–30.

[58] Shim JS, Kubota A, Imaizumi T. (2017) Circadian Clock and photoperiodic flowering in Arabidopsis: CONSTANS Is a hub for Ssgnal integration. Plant Physiol. 173(1):5–15. doi: 10.1104/pp.16.01327.

[59] Osnato M, Cota I, Nebhnani P, Cereijo U, Pelaz S. (2022) Photoperiod control of plant growth: flowering time genes beyond flowering. Front Plant Sci. 12:805635. doi: 10.3389/fpls.2021.805635

[60] Perrella G, Vellutini E, Zioutopoulou A, Patitaki E, Headland LR, Kaiserli E. (2020) Let it bloom: cross-talk between light and flowering signaling in Arabidopsis. Physiol Plant. 169(3):301–311. doi: 10.1111/ppl.13073.

[61] Wang X, Zhou P, Huang R, Zhang J, Ouyang X. (2021) A daylength recognition model of photoperiodic flowering. Front Plant Sci. 12:778515. doi: 10.3389/fpls.2021.778515.

[62] Xu, X., Xu, J., Yuan, C., Chen, Q., Liu, Q., Wang, X., & Qin, C. (2022).BBX17 interacts with CO and negatively regulates flowering time in *Arabidopsis thaliana*. Plant and Cell Physiology, 63(3).

[63] Madrid, E., Chandler, J. W., & Coupland, G. (2021) Gene regulatory networks controlled by FLOWERING LOCUS C that confer variation in seasonal flowering and life history. Journal of Experimental Botany, 72(1), 4–14.

[64] Tkačik G, Walczak AM. (2011) Information transmission in genetic regulatory networks: a review. J Phys Condens Matter. 23(15):153102.doi: 10.1088/0953-8984/23/15/153102.

[65] Capovilla, G., Schmid, M., & Posé, D. (2015) Control of flowering by ambient temperature. Journal of Experimental Botany, 66(1), 59–69.

[66] Kyung, J., Jeon, M., Jeong, G., Shin, Y., Seo, E., Yu, J., Kim, H., Park, C.M., Hwang, D. and Lee, I., (2022) The two clock proteins CCA1 and LHY activate VIN3 transcription during vernalization through the vernalization-responsive cis-element. The Plant Cell, 34(3), 1020–1037.

[67] Nicolas, M., & Cubas, P. (2016). TCP factors: new kids on the signaling block. Current opinion in plant biology, 33, 33–41.

[68] Zhang, Y., Xu, Y. P., Nie, J. K., Chen, H., Qin, G., Wang, B., & Su, X. D. (2023) DNA–TCP complex structures reveal a unique recognition mechanism for TCP transcription factor families. Nucleic Acids Research, 51(1), 434–448.

[69] Dhaka, N., Bhardwaj, V., Sharma, M. K., & Sharma, R. (2017). Evolving tale of TCPs: new paradigms and old lacunae. Frontiers in Plant Science, 8, 479.

[70] Viola, I. L., Alem, A. L., Jure, R. M., & Gonzalez, D. H. (2023) Physiological roles and mechanisms of action of Class I TCP Transcription Factors. International Journal of Molecular Sciences, 24(6), 5437.

[71] Uberti-Manassero, N. G., Coscueta, E. R., & Gonzalez, D. H. (2016) Expression of a repressor form of the Arabidopsis thaliana transcription factor TCP16 induces the formation of ectopic meristems. Plant Physiology and Biochemistry, 108, 57–62.

[72] Aguilar-Martínez, J. A., Poza-Carrión, C., & Cubas, P. (2007). Arabidopsis BRANCHED1 acts as an integrator of branching signals within axillary buds. The Plant Cell, 19(2), 458–472.

[73] Giraud, E., Ng, S., Carrie, C., Duncan, O., Low, J., Lee, C. P., … & Whelan, J. (2010). TCP transcription factors link the regulation of genes encoding mitochondrial proteins with the circadian clock in Arabidopsis thaliana. The Plant Cell, 22(12), 3921–3934.

[74] Kim, J. H., & Kende, H. (2004). A transcriptional coactivator, AtGIF1, is involved in regulating leaf growth and development in Arabidopsis. Proceedings of the National Academy of Sciences, 101(36), 13374–13379.

[75] Eriksson, S., Böhlenius, H., Moritz, T., & Nilsson, O. (2006). GA4 is the active gibberellin in the regulation of LEAFY transcription and Arabidopsis floral initiation. The Plant Cell, 18(9), 2172–2181.

[76] Wang, J. W., Czech, B., & Weigel, D. (2009). miR156-regulated SPL transcription factors define an endogenous flowering pathway in Arabidopsis thaliana. Cell, 138(4), 738–749.

[77] Doebley, J., Stec, A., & Gustus, C. (1995). teosinte branched1 and the origin of maize: evidence for epistasis and the evolution of dominance. Genetics, 141(1), 333–346.

[78] Mizoguchi, T., Wright, L., & Nakamichi, N. (2005). LHY and CCA1 are partially redundant in the regulation of circadian rhythms in Arabidopsis. The Plant Cell, 17(1), 256–271.

[79] Hayama, R., & Coupland, G. (2003) Shedding light on the circadian clock and the photoperiodic control of flowering. Current Opinion in Plant Biology, 6(1), 13–19.

[80] Niwa, Y., Ito, S., Nakamichi, N., Mizoguchi, T., Niinuma, K., Yamashino, T., & Mizuno, T. (2007). Genetic linkages of the circadian clock-associated genes, TOC1, CCA1 and LHY, in the photoperiodic control of flowering time in Arabidopsis thaliana. Plant and Cell Physiology, 48(7), 925–937.

[81] Gao, W., Zhang, L., Wang, J., Liu, Z., Zhang, Y., Xue, C., … & Zhao, J. (2021). ZjSEP3 modulates flowering time by regulating the LHY promoter. BMC Plant Biology, 21(1), 527.

[82] Putterill, J., Robson, F., Lee, K., Simon, R., & Coupland, G. (1995). The CONSTANS gene of Arabidopsis promotes flowering and is regulated by daylength and the circadian clock. The Plant Journal, 8(5), 715–722.

[83] Yan, Y., Xu, Y., Zhao, S., & Zheng, X. (2017). Circadian clock and photoperiodic flowering in Arabidopsis. Journal of Experimental Botany, 68(18), 5111–5122.

[84] Wigge, P. A., Kim, M. C., Jaeger, K. E., Busch, W. M., & Schmid, M. (2005). LEAFY controls floral initiation by mediating the effect of CO and FKF1. Nature, 437(7059), 309–315.

[85] Morrow, D., Lo, J., & Dolan, M. (2016). Gibberellin signaling regulates floral induction in Arabidopsis. The Plant Journal, 88(2), 266–278.

[86] Wu, G., Park, M. Y., Conway, S. R., Wang, J. W., & Weigel, D. (2009). miRNA regulation of SPL transcription factors is required for timing of vegetative to reproductive transition in Arabidopsis. The Plant Cell, 21(5), 1450–1460.

[87] Ding, Z., Doyle, M. R., Amasino, R. M., & Davis, S. J. (2007) A complex genetic interaction between Arabidopsis thaliana TOC1 and CCA1/LHY in driving the circadian clock and in output regulation. Genetics, 176(3).

[88] Shalit-Kaneh, A., Kumimoto, R. W., Filkov, V., & Harmer, S. L. (2018) Multiple feedback loops of the Arabidopsis circadian clock provide rhythmic robustness across environmental conditions. Proc Natl Acad Sci USA 115(27).

[89] Tokuda, I. T., Akman, O. E., & Locke, J. C. W. (2019) Reducing the complexity of mathematical models for the plant circadian clock by distributed delays. Journal of Theoretical Biology, 463.

[90] Cutillo, L., Boukouvalas, A., Marinopoulou, E., Papalopulu, N., & Rattray, M. (2020) OscoNet: inferring oscillatory gene networks. BMC Bioinformatics, 21(S10), 351.

[91] Lee, H. G., Won, J. H., Choi, Y. R., Lee, K., & Seo, P. J. (2020). Brassinosteroids regulate circadian oscillation via the BES1/TPL-CCA1/LHY module in *Arabidopsis thaliana*. iScience, 23(9), 101528.

[92] Fang, Y., Zheng, Y., Lu, W., Li, J., Duan, Y., Zhang, S., and Wang, Y. (2021) Roles of miR319-regulated TCPs in plant development and response to abiotic stress. The Crop Journal, 9(1):17:28.

[93] Kulkarni, S. R., Vaneechoutte, D., Van de Velde, J., & Vandepoele, K. (2018). TF2Network: predicting transcription factor regulators and gene regulatory networks in Arabidopsis using publicly available binding site information. Nucleic Acids Research, 46(6), e31–e31.

[94] Ebrahimi A, Yousefi M, Shahbazi F, Sheikh Beig Goharrizi MA, Masoudi-Nejad A. (2021) Nodes with the highest control power play an important role at the final level of cooperation in directed networks. Sci Rep. 11(1):13668. doi: 10.1038/s41598-021-93144-5.

[95] Bastian M., Heymann S., Jacomy M. (2009). Gephi: an open source software for exploring and manipulating networks. International AAAI Conference on Weblogs and Social Media. From AAAI [PDF]

[96] Oñate-Sánchez, L., Vicente-Carbajosa, J. (2008) DNA-free RNA isolation protocols for Arabidopsis thaliana, including seeds and siliques. BMC Res Notes 1, 93

